# Crowdsourced mapping extends the target space of kinase inhibitors

**DOI:** 10.1101/2019.12.31.891812

**Authors:** Anna Cichonska, Balaguru Ravikumar, Robert J Allaway, Sungjoon Park, Fangping Wan, Olexandr Isayev, Shuya Li, Michael Mason, Andrew Lamb, Ziaurrehman Tanoli, Minji Jeon, Sunkyu Kim, Mariya Popova, Stephen Capuzzi, Jianyang Zeng, Kristen Dang, Gregory Koytiger, Jaewoo Kang, Carrow I. Wells, Timothy M. Willson, The IDG-DREAM Drug-Kinase Binding Prediction Challenge Consortium, Tudor I. Oprea, Avner Schlessinger, David H. Drewry, Gustavo Stolovitzky, Krister Wennerberg, Justin Guinney, Tero Aittokallio

## Abstract

Despite decades of intensive search for compounds that modulate the activity of particular targets, there are currently small-molecules available only for a small proportion of the human proteome. Effective approaches are therefore required to map the massive space of unexplored compound-target interactions for novel and potent activities. Here, we carried out a crowdsourced benchmarking of predictive models for kinase inhibitor potencies across multiple kinase families using unpublished bioactivity data. The top-performing predictions were based on kernel learning, gradient boosting and deep learning, and their ensemble resulted in predictive accuracy exceeding that of kinase activity assays. We then made new experiments based on the model predictions, which further improved the accuracy of experimental mapping efforts and identified unexpected potencies even for under-studied kinases. The open-source algorithms together with the novel bioactivities between 95 compounds and 295 kinases provide a resource for benchmarking new prediction algorithms and for extending the druggable kinome.

## Introduction

Only 11% of the human proteome can be currently targeted by small molecules or drugs, whereas one in 3 proteins remains under-studied.^1^ Furthermore, despite many years of target-based drug discovery, chemical agents inhibiting single protein targets are still rare.^2^ Most approved drugs have multiple targets, suggesting their therapeutic efficacy as well as adverse side-effects originate from polypharmacological effects.^3^ Systematic mapping of the target binding profiles is therefore critical not only to explore the therapeutic potential of promiscuous agents, but also to better predict and manage their possible adverse effects prior to further development and clinical trials (i.e., speeding-up and de-risking the drug development process). Comprehensive understanding of the pharmacological effects of approved drugs could uncover novel off-target potencies to extend their therapeutic application area (via off-label use or repurposing). However, the massive size of the chemical universe makes the complete experimental mapping of compound-target activities infeasible, even with automated high-throughput profiling assays.

To accelerate the mapping efforts, we implemented the IDG-DREAM Drug-Kinase Binding Prediction Challenge, a crowd-sourced competition that evaluated the power of machine learning (ML) models as a systematic and cost-effective means for predicting novel compound-target potencies that warrant experimental evaluation (i.e., target prioritization). The Challenge focused on kinase inhibitors, since kinases are tractable in drug development and play a role in a wide range of diseases, such as cardiovascular disorders and cancers. However, protein kinase domains share structural and sequence similarity, and most kinase inhibitors bind to conserved ATP-binding pockets, which leads to prevalent target promiscuity and polypharmacological effects.^4–7^ Such promiscuity requires effective target deconvolution approaches, including ML approaches, that can leverage the information extracted from similar kinases and compounds to predict the activity of so far unexplored interactions.^8,9^

The Challenge was implemented in a screening-based, pre-competitive drug discovery project in collaboration with the NIH-supported Illuminating the Druggable Genome (IDG) program (https://commonfund.nih.gov/idg), with the common aim to establish kinome-wide target profiles of small-molecule agents, and thereby to extend the druggability of the human kinome space by providing activity information on under-studied proteins. The specific questions this Challenge sought to address were: (i) What are the best computational modelling approaches for predicting quantitative compound-target activity profiles?; (ii) What are the optimal molecular and chemical descriptors for maximal prediction accuracy?; and (iii) What are the most predictive bioactivity assays and publicly available resources? The Challenge attracted 212 active participants, and a total of 268 predictions were scored, covering a wide range of ML approaches, including linear regularized regression, deep and kernel learning algorithms and gradient boosting decision trees. Here, we describe the benchmarking results from the Challenge, and the use of top-performing prediction models for identifying novel kinase inhibitor activities.

## Results

### Challenge implementation

To develop their predictive models, the participants had access to a wide variety of bioactivity data for model training and cross-validation through open databases such as ChEMBL^10^, BindingDB^11^ and IDG Pharos^12^ (Fig. 1). For training data collection, integration, management and harmonization, the Challenge made use of an open-data platform, DrugTargetCommons (DTC).^13^ DTC is a community platform that facilitates the annotation and curation of bioactivity data, and provides a comprehensive and standardized interface to retrieve compound-target profiles and related information to support predictive modelling (Suppl. Fig. 1). The Challenge infrastructure was built on the Synapse collaborative science platform^14^, which supported receiving, validating and scoring of the teams’ predictions as well as long-term management of the test bioactivity data and submitted Challenge models as a benchmarking resource (Fig. 1).

**Figure 1.**
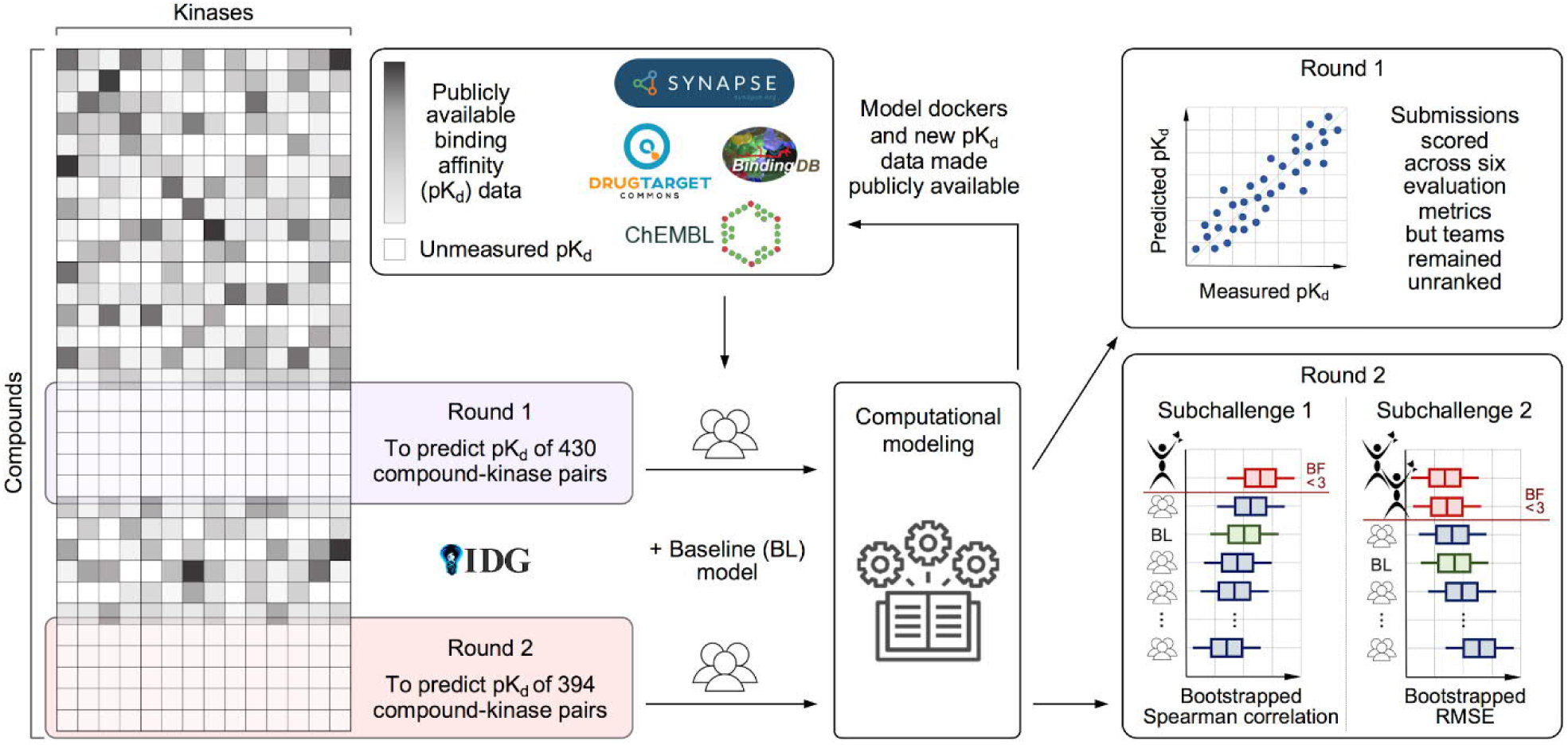
Overview of the IDG-DREAM Drug-Kinase Binding prediction Challenge. The participants had access to publicly available target profiling training data, and the predictions were then validated in two unpublished and blinded test data sets profiled by the Illuminating the Druggable Genome (IDG) program. The heatmap on the left is for illustrative purposes only (see Suppl. Fig. 2 for the actual test data matrices, and Suppl. Fig. 3 for the Challenge timeline).

### Challenge test data sets

Evaluation of the model predictions was based on unpublished target binding data generated by the IDG Kinase Data and Resource Generation Center, conducted over a series of “rounds” based on availability of validation datasets (Suppl. Fig. 3). Generation of the test data for Round 1 was based on a single-dose kinome scan of a library of multi-targeted compounds.^6,15^ This was followed by a dose-response determination of the dissociation constant (K_d_) values for 430 compound-kinase pairs between 70 inhibitors and 199 kinases that were not available in the public domain (see Methods). An additional set of completely new K_d_ data was generated for Round 2, consisting of 394 multi-dose assays between 25 inhibitors and 207 kinases with single-dose inhibition >80%. Together, these 824 K_d_ assays in the two Rounds spanned a total of 95 compounds and 295 kinases (Fig. 2a-b), consisting of promiscuous compounds targeting multiple kinases at low concentrations, compounds with narrow target profiles, as well as compounds with no potent targets among the tested kinases (Suppl. Fig. 2).

**Figure 2.**
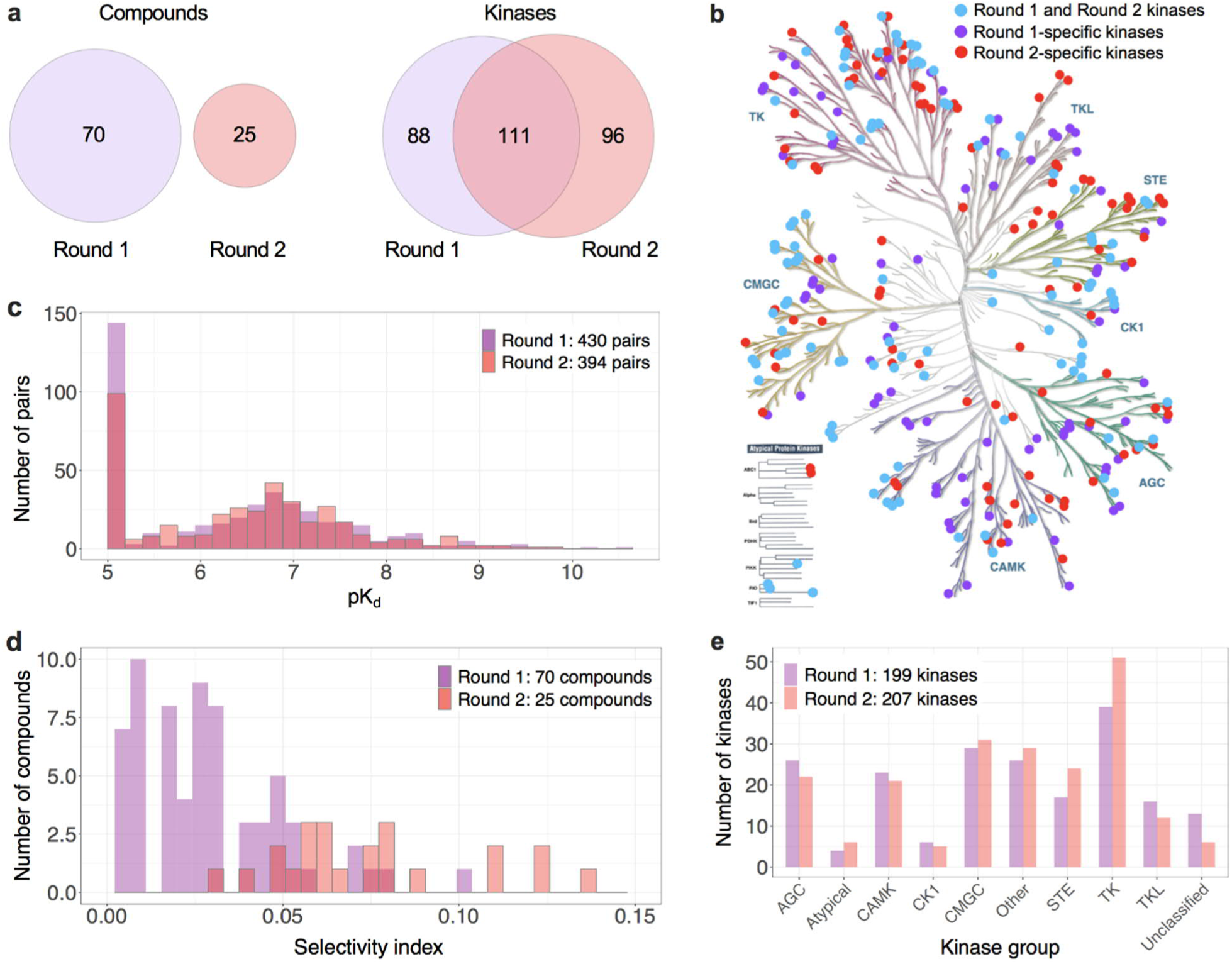
Challenge test datasets. (a) The overlap between Round 1 and Round 2 test compounds and kinases, and their distributions in the kinome tree (b), and across kinase groups (e). (c) The quantitative dissociation constant (K_d_) of compound-kinase activities was measured in dose-response assays (see Methods), presented in the logarithmic scale as pK_d_ = −log_10_(K_d_). The higher the pK_d_ value, the higher the inhibitory ability of a compound against a protein kinase (Suppl. Fig. 2 lists the compounds and kinases in Round 1 and Round 2). (d) The selectivity index for compounds was calculated based on the single-dose activity assay (at 1 µM of compound) across full compound-kinase matrices before the Challenge. The kinome tree figure was created with KinMap, reproduced courtesy of Cell Signaling Technology, Inc.

Round 1 enabled the teams to carry out initial testing of various model classes and data resources, whereas Round 2, implemented 6 months later, was used to score the final prediction models and to select the top-performing teams. Round 1 test data remained blinded in Round 2. Round 1 and 2 test data had very similar K_d_ distributions (Fig. 2c), which provided comparable binding affinity outcome data to monitor the improvements made by the teams between the two rounds. Compounds in the test sets were mutually exclusive between rounds (Fig. 2a), with Round 2 including less selective compounds with broader target profiles (Fig. 2d), and therefore fewer inactive compound-target pairs (pK_d_=5). Round 1 and 2 kinase targets were partly overlapping, and covered all major kinase families and groups (Fig. 2b,e). Taken together, these two test datasets provided a standardized and sufficiently large quantitative bioactivity resource to evaluate the accuracy of predicting on- and off-target activities.

### Predictive performance of the models

The competition challenged the participants to predict blinded K_d_ profiles between 95 compounds and 295 kinases. Since a goal of this Challenge was to encourage algorithm development that would exceed state-of-the-art, we selected as “base-line model” a recently published and experimentally-validated kernel regression approach for compound-kinase activity prediction^16^. The performance of the predictions improved from Round 1 to Round 2 submissions as measured by Spearman correlation (two-sample Wilcoxon test, p<0.005; Fig. 3a) and Root Mean Square Error (RMSE, p<10^−6^; Fig. 3c). Comparison against the baseline model indicated that the Round 2 dataset was marginally easier to predict (Suppl. Fig. 4), partly due to a smaller proportion of inactive pairs in Round 2 (pK_d_ = 5, Fig. 2c). To take into account this shift, we compared the submissions against a set of random predictions. Using Spearman correlation, we observed that 48% of the submission were better than random in Round 1, compared to 61% in Round 2 (Fig 3b). Using RMSE, 71% of the submissions in Round 1 were better than random, compared to 76% in Round 2 (Fig 3d).

**Figure 3.**
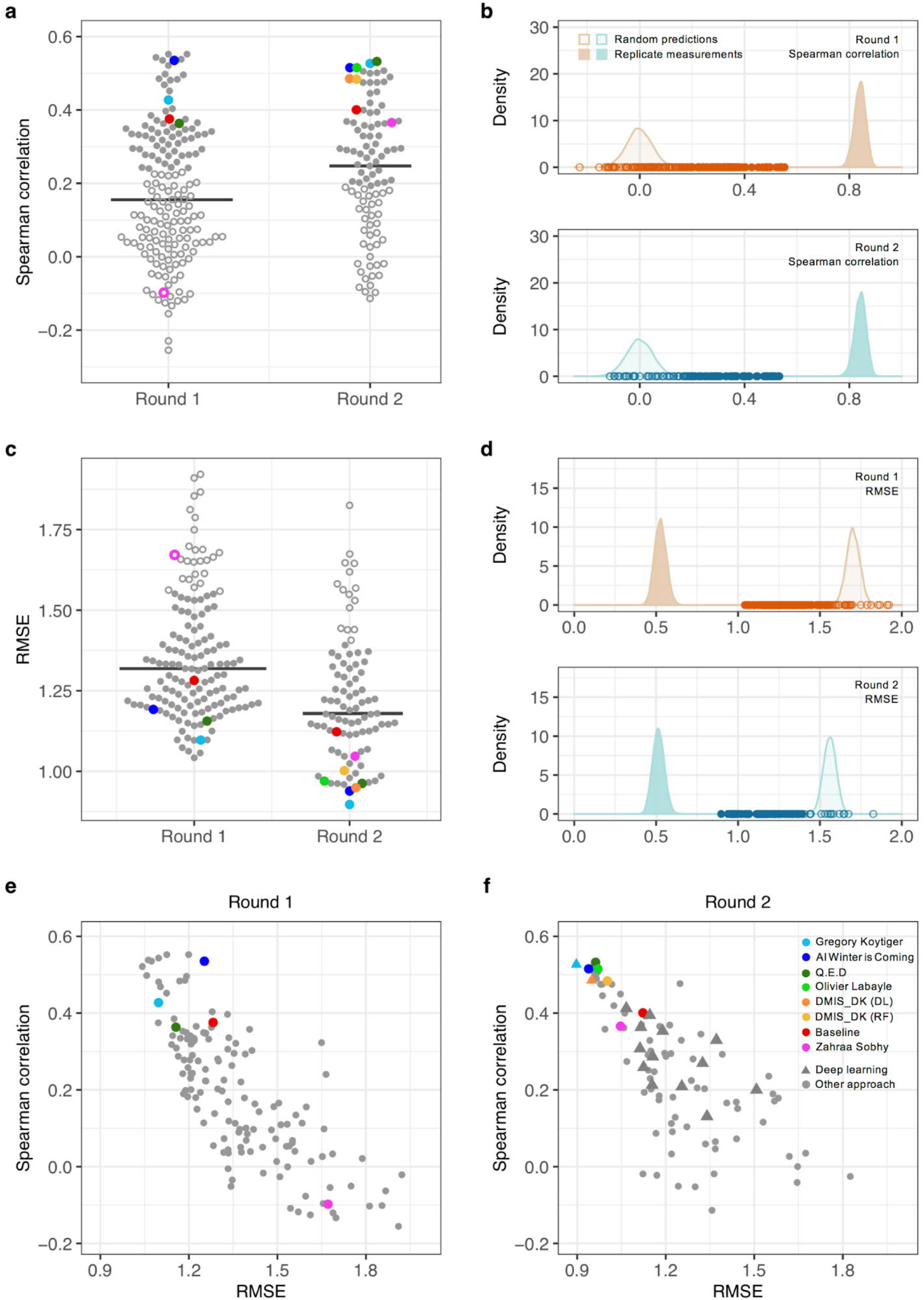
Overall performance of the submissions. (a, c) Performance of the submissions in terms of the two winning metrics in Round 1 (*n* = 169 submissions) and Round 2 (*n* = 99). The colors mark the baseline model and the top-performing participants in Round 2. The empty circles mark the submissions that did not differ from random predictions. The baseline model^16^ remained the same in both of the rounds. (b, d) Distribution of the random predictions (based on 10,000 permuted pK_d_ values) and replicate distributions (based on 10,000 subsamples with replacement of overlapping pK_d_ pairs between two large-scale target activity profiling studies^4,5^) in Round 1 (top panel) and Round 2 (bottom). The points correspond to the individual submissions. (e, f) Relationship of the two winning metrics across the submissions in Round 1 and Round 2. The shape indicates submissions based on deep learning in Round 2 (f). For instance, team DMIS_DK submitted predictions based both on random forest (RF) and deep learning (DL) algorithms in Round 2, where the latter showed slightly better accuracy (triangle).

The 20 teams that participated in both rounds improved their K_d_ predictions (p<0.05 and p<0.001 for Spearman and RMSE, paired Wilcoxon signed-rank test), but when comparing against the baseline model, the overall improvements became insignificant (p>0.05). However, there were individual teams (like Zahraa Sobhy) that were able to improve their predictions considerably between the two rounds. The practical upper bound of the model predictions was defined based on experimental replicates of K_d_ measurements (Fig. 3b,d). The predictive accuracy of the top-performing models in Round 2 was relatively high based on both of the winning metrics, Spearman correlation for rank predictions and RMSE for activity predictions; these metrics showed less correlated performance over the less-accurate models in Round 2 (Fig. 3f). The tie breaking metric, averaged area under the curve (AUC), provided complementary information on prediction accuracy when compared to RMSE but not to Spearman correlation (Suppl. Fig. 5). Overall, models based on deep learning algorithms did not perform better than the other leaning algorithms submitted in Round 2 (Fig. 3f).

### Analysis of the top-performing models

The top-performing models were selected in Round 2 based on 394 pK_d_ predictions between 25 compounds and 207 kinases. Only those participants who submitted their Dockerized models, method write-ups, and method surveys were qualified to win the two sub-challenges. To select the top-performers, we conducted a bootstrap analysis of each participant’s best submission, and then calculated a Bayes factor (K) relative to the bootstrapped overall best submission for each winning metric (Suppl. Fig. 6). Considering Spearman correlation, the top-performer was team Q.E.D (K<3; Fig. 4a). For the RMSE metric, the top-performing teams were AI Winter is Coming (AIWIC) and DMIS_DK (K<3; Suppl. Fig. 6), with AIWIC having a marginally better tie-breaking metric (average AUC of 0.773; Fig. 4b). Only two non-qualifying participants (Gregory Koytiger and Olivier Labayle) showed comparable performance. Overall, these five teams performed the best when considering the 54 teams in Round 2 (Suppl. Fig. 7).

**Figure 4.**
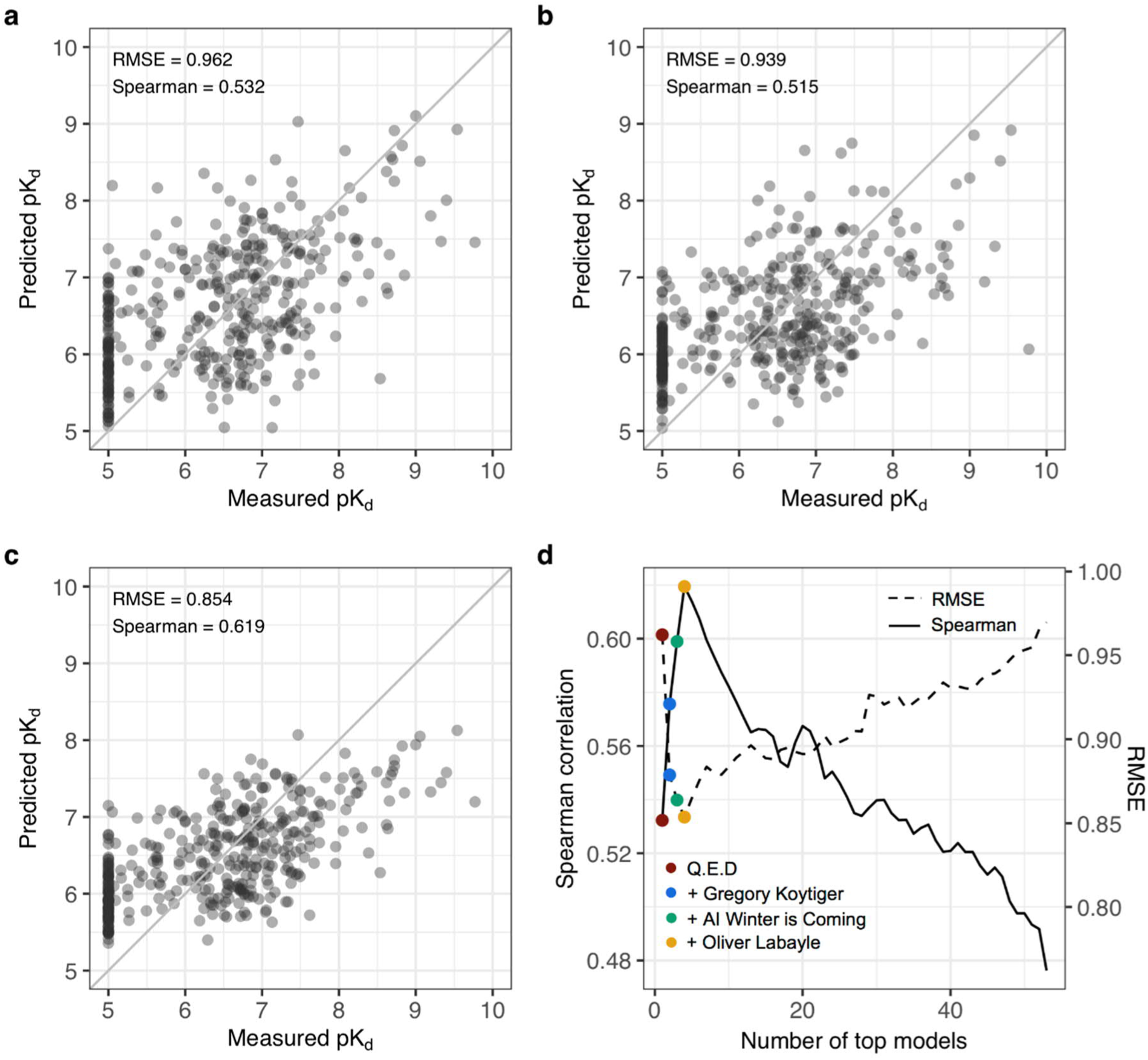
Top-performing models and their ensemble combination. (a) Spearman correlation sub-challenge top-performer in Round 2 (Q.E.D). (b) RMSE sub-challenge top-performer in Round 2 (AI Winter is Coming). Points correspond to 394 pairs between 25 compounds and 207 kinases. (c) Ensemble model that combines the top four models selected based on their Spearman correlation in Round 2. (d) The mean aggregation ensemble model was constructed by adding an increasing number of top-performing models based on their Spearman correlation (the solid curve), until the ensemble correlation dropped below 0.45. The peak performance was reached after aggregating four teams (marked in the legend; see Suppl. Fig. 8 for names of all the teams). The right-hand y-axis and the dotted curve shows the RMSE of the ensemble model as a function of an increasing number of top-performing models.

Notably, the top-performing models were based on various ML approaches, including deep learning, graph convolutional networks, gradient boosting decision trees, kernel learning and regularized regression (Table 1). To study whether combining predictions from multiple ML approaches could improve prediction accuracy, we constructed an ensemble model by simple mean aggregation of an increasing number of top-performing models in Round 2. The combination of the four best performing models resulted in the peak Spearman correlation (Fig. 4c), demonstrating complementary value of these predictions. After adding more models, the ensemble prediction accuracy decreased rapidly, measured by Spearman correlation and RMSE (Fig. 4d). However, an ensemble prediction from a total of 21 best teams had a significantly better correlation than the best single model alone (K>5; Suppl. Fig. 8). This suggests that the combination of various ML approaches using an ensemble model leads to accurate and robust predictions of kinase inhibitor potencies across multiple kinase families.

**Table 1.** Model classes and training data of the Round 2 top-performing teams and the baseline model^16^. Even if the teams chose to combine predictions from multiple models, they had to submit only one prediction per compound-target pair for scoring against the measured activities.

| **Team** | **Algorithm type** | **Algorithm name** | **Combined models** | **Training strategy** |
| --- | --- | --- | --- | --- |
| *DMIS_DK* | Deep learning | Graph Neural Networks | 12 | Train test split |
| *AI Winter is Coming* | Gradient boosting decision trees | Xgboost | 5 per target | K-fold nested cross validation, boosting |
| *Q.E.D* | Kernel learning | CGKronRLS | 440 | Boosting |
| *Gregory Koytiger* | Deep learning | Not applicable | 6 | Fixed hold out |
| *Olivier Labayle* | Ridge regression | Not applicable | Not applicable | K-fold cross validation |
| *Baseline* | Kernel learning | CGKronRLS | 1 | K-fold nested cross validation |

| **Team** | **Training data sources** | **Compound-protein pairs** | **Bioactivity types** | **Protein features** | **Chemical features** |
| --- | --- | --- | --- | --- | --- |
| *DMIS_DK* | DrugTargetCommons, BindingDB | 953521 | K<sub>d</sub>, K<sub>i</sub>, IC<sub>50</sub> | None | 2D molecular graphs |
| *AI Winter is Coming* | DrugTargetCommons, ChEMBL | 600000 | K<sub>d</sub>, K<sub>i</sub>, IC<sub>50</sub>, EC<sub>50</sub> | None | ECFP5, ECFP7, ECFP9, ECFP11 |
| *Q.E.D* | DrugTargetCommons, ChEMBL, Uniprot | 60462 | K<sub>d</sub>, K<sub>i</sub>, EC<sub>50</sub> | Amino acid sequences | ECFP4, ECFP6 |
| *Gregory Koytiger* | ChEMBL | 250000 | K<sub>d</sub>, K<sub>i</sub>, IC<sub>50</sub> | None | None |
| *Olivier Labayle* | DrugTargetCommons, ChEMBL, Uniprot | 18200 | K<sub>d</sub> | K-mer counting | ECFP |
| *Baseline* | DrugTargetCommons | 44186 | K<sub>d</sub> | Amino acid sequences | Path-based fingerprints |

### Comparison against single-dose activity

We next investigated how well the top-performing ML models compare against single-dose activity assays in terms of reducing the number of false positives and negatives when selecting most potent compound-target activities for more detailed, multi-dose K_d_ profiling. For this classification task, we defined the ground truth activity classes based on the measured K_d_ potencies, which provide a more practical prediction outcome, compared to the rank correlation analyses that already demonstrated predictive rankings with the top-performing models (Fig. 4). Using the activity cut-off of measured pK_d_ = 6 and an single-dose inhibition cut-off of 80%, similar to previous studies,^6,15,17^ the positive predictive value (PPV) and the false discovery rate (FDR) of the single-dose assay were PPV = 0.66 and FDR = 0.44 in the Round 2 dataset. When using the mean aggregation ensemble of the predicted pK_d_ values from the top-performing models and the same cut-off of pK_d_ > 6 for both the predicted and measured activities, we observed an improved precision of PPV = 0.76 and FDR = 0.24.

To further repeat the activity classification with multiple cut-off levels, we ranked the Round 2 pairs both using the model-predicted pK_d_ values and the measured single-dose inhibition assay values, and then compared these rankings against the measured dose-response assay (here, pK_d_ > 6 indicates positive activity class). The ROC analyses demonstrated an improved activity classification accuracy using the mean ensemble of the top-performing models (Fig. 5a), especially when focusing on the most potent compound-target activities with the highest specificity. This improvement in both sensitivity and specificity was achieved without making any additional activity measurements, and it became even more pronounced with the precision-recall analysis, which showed that the precision of the prediction models remained above PPV=75% level even when the recall (sensitivity) level exceeded 75% (Fig. 5b). As expected, the prediction accuracy decreased when using a more stringent activity cut-off of pK_d_ > 7 (Suppl. Fig. 9), since these rare extreme activities are more challenging to predict.

**Figure 5.**
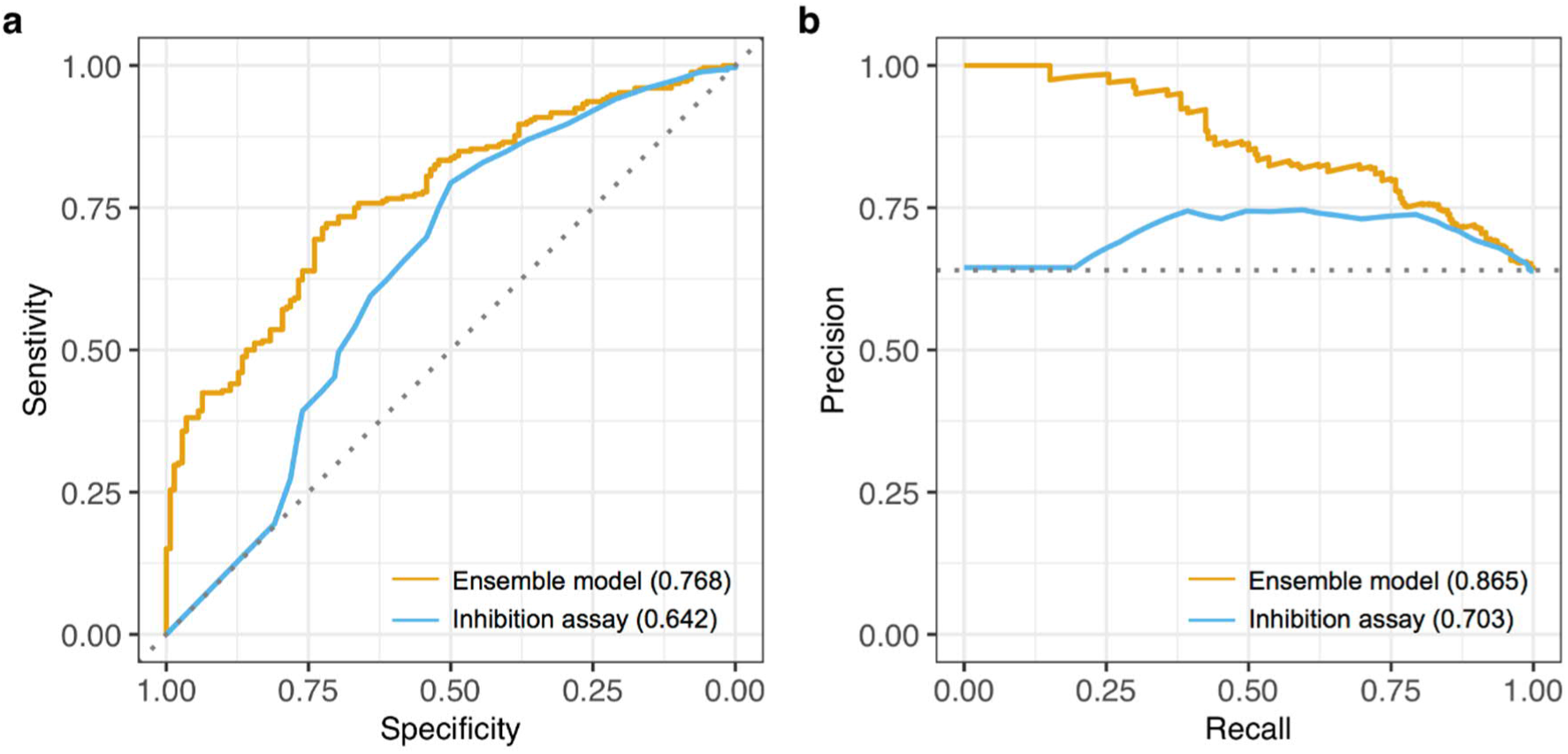
Top-performing model predictions compared against single-dose assays. (a) Receiver operating characteristic (ROC) curves when ranking the 394 compound-kinase pairs from Round 2 using both the ensemble of the top-performing models (average predicted pK_d_ from Q.E.D, DMIS_DK and AI Winter is Coming) and the experimental single-dose inhibition assays (the pairs with higher inhibition% are ranked first). The true positive activity class includes pairs with measured pK_d_ > 6. The area under the ROC curve values are shown in the parentheses and the diagonal dotted line shows the random prediction accuracy of AU-ROC=0.50. (b) Precision-recall (PR) curves for the same activity classification analysis as shown in panel a. The area under the PR curve values are shown in parentheses and the horizontal dotted line indicates the random classifier precision of 0.64. *Note*: Round 2 K_d_ measurements were pre-selected to include mostly those pairs with single-dose inhibition>80%, which makes Round 2 pairs optimal for systematic analysis of false positive predictions, and hence sensitivity (recall) and PPV (precision). However, these 394 pairs pre-selected for K_d_ profiling were less optimal for a comprehensive analysis of false negative predictions, and the evaluation of specificity.

### Model-based target predictions

To further analyse both the sensitivity and specificity of the model predictions, we experimentally profiled 81 additional pairs, which were not part of Round 1 or 2 datasets, selected based on the pK_d_ predictions from the three top-performing models. These follow-up experiments were carried out in an unbiased manner, regardless of the compound classes, kinase families, or inhibition levels, to investigate the accuracy of predictive models to identify potent inhibitors of kinases with less than 80% single-dose inhibition; this activity cut-off is often used when selecting pairs for multi-dose K_d_ testing^6,15,17^, but it may miss the more challenging compound-kinase pairs with lower single-dose inhibition. Most of the measured pK_d_ values of these 81 pairs were distributed as expected, according to the expected single-dose inhibition function (Fig. 6a, black trace). However, our model-based approach also identified unexpected activities (pK_d_ > 6), that could not be predicted based on the inhibition assay only; those with pK_d_ > 7 are discussed below.

**Figure 6.**
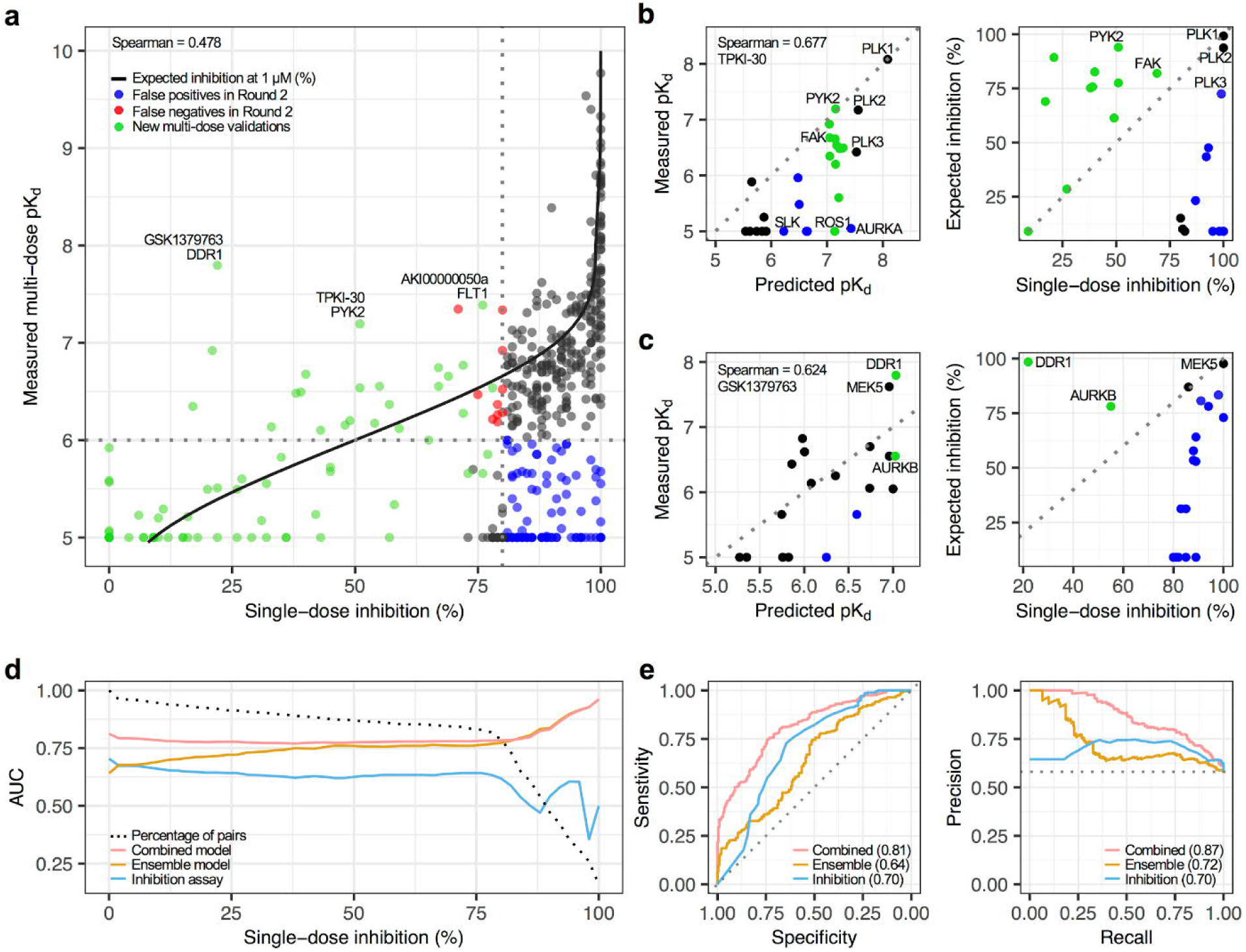
Machine learning-based target predictions. (a) Comparison of single-dose inhibition assay (at 1 µM) against multi-dose K_d_ assay activities across 475 compound-target pairs (394 Round 2 pairs and 81 additionally profiled pairs). The red points indicate false negatives and blue points false positives when using cut-offs of pK_d_ = 6 and inhibition=80% among the Round 2 pairs (including 75 pairs with inhibition>80% but that showed no activity in the dose-response assays, i.e, pK_d_ = 5). The green points indicate the new experiments carried out solely based on the model predictions, regardless of inhibition levels. The black trace indicates the expected %inhibition rate based on measured pK_d_’s, estimated using the maximum ligand concentration of 1 µM both for the single-dose and dose-response assays (see Methods). (b) Multi-dose (left) and single-dose (right) assays for kinases tested with TPKI-30. Green points indicate the new experimental validations based on model predictions, whereas black points come from Round 2 data. Blue points indicate false positive predictions based either on predictive models or single-dose testing. (c) Multi-dose (left) and single-dose (right) assays for kinases tested with GSK1379763. The color-cording is the same as in panel b. (d) Predictive accuracy of the ensemble of top-performing models (average predicted pK_d_) and single-dose assay (at 1 µM), when classifying subsets of the 475 pairs into those with measured pK_d_ less or higher than 6. The y-axis indicates the area under the receiver operating characteristic curve (AUC) as a function of the single-dose inhibition% levels, x-axis indicates the pairs with inhibition>x%, and the dotted black curve the percentage of all pairs that passed that activity threshold. The combined model trace corresponds to the average of measured and expected inhibition values, where the latter was calculated based on the mean ensemble of the top-performing model pK_d_ predictions (Q.E.D., DMIS_DK and AI Winter is Coming). (e) Receiver operating characteristic (ROC) curves (left) and precision-recall (PR) curves (right), when ranking all the 475 pairs using the top-performing model-predicted pK_d_ values and the measured single-dose inhibition assays, or using their combination. The AUC values are shown in the parentheses. The diagonal dotted line indicates the random prediction accuracy of AU-ROC=0.50 (left), and the horizontal dotted line indicates the random classifier precision of 0.58 (right).

As an example of a potent activity missed by the single-dose assays, the top-performing models predicted PYK2 (PTK2B) as a high-affinity target of a PLK inhibitor TPKI-30 (Fig. 6a). The new multi-dose pK_d_ measurements validated that TPKI-30 indeed has an activity against PYK2 close to its potency towards PLK2 (Fig. 6b, left panel). Neither PYK2 or FAK would have been predicted as potent targets based on the single-dose testing alone, which led to multiple false negatives (Fig. 6b, right panel). In general, the single dose-testing had a relatively low predictivity of actual TPKI-30 potencies, since kinases other than PLKs with high single-dose activity were confirmed as non-potent targets based on dose-response K_d_ testing, resulting in many false positives. In contrast, the top-performing model predictions turned out to be relatively accurate, except for a few receptor tyrosine kinases (Fig. 6b, left panel).

Another unexpected target activity was predicted for GSK1379763 that also showed high potency against DDR1 based on the subsequent K_d_ assays, exceeding that of the AURKB (Fig. 6c, left panel). The single-dose testing suggested that this compound would have potency neither against DDR1 or AURKB (Fig. 6c, right panel), whereas the multi-dose assays confirmed potency towards DDR1 at a similar level as the Round 2 highest affinity target MEK5 (MAP2K5). Whereas DDR1 has been studied extensively^18^, there are only a few activity data points currently available for the other high-affinity target, PYK2, suggesting that the prediction models can identify potent inhibitors even for under-studied kinases that would have been missed when using the single-dose assays alone. In contrast, the third predicted activity between AKI00000050a and FLT1 could have been identified based on its relatively high single-dose activity, even if less than 80% (Fig. 6a).

Surprisingly, the single-dose assays and model-based pK_d_ predictions were weakly correlated (Suppl. Fig. 10, Spearman correlation 0.24), and they showed opposite trends for K_d_ prediction accuracy when increasing the inhibition cut-off level (Fig. 6d). To combine these two activity estimators, we calculated for each compound-kinase pair an average of its measured and expected inhibition values based on the single-dose assay and the top-performing models, respectively. This combined predictor showed improved activity classifications beyond that of the model predictions alone, across various inhibition levels, and identified an extended number of potent compound-target interactions at lower single-dose activity, compared to the standard 80% cut-off (Fig. 6d, dotted line). The combined model improved both the sensitivity and specificity of the pK_d_ predictions among all the 475 pairs (Fig. 6e, left panel), and especially the precision of the top-activity predictions that are prioritized for further validation (Fig. 6e, right panel).

## Discussion

While experimental mapping of target activities is critical for understanding compounds’ mode of action (MoA), biochemical target activity profiling experiments are both time consuming and costly. The enormous size of the chemical universe, spanned by up to 10^20^ molecules with potential pharmacological properties,^19,20^ makes the experimental bioactivity mapping of the full compound and target space quickly infeasible in practice. The IDG-DREAM Drug Kinase Binding Prediction Challenge was designed to benchmark algorithms capable of predicting and prioritizing compound-target activities, and therefore to guide data-driven decision making and reduce the high failure rates. The model-guided approach has the potential to help both phenotype-based drug discovery (e.g. mapping of the active target space of lead compounds), and target-based drug discovery (e.g. identification of candidate compounds that selectively inhibit a particular disease-related target). As an example, the top-performing models led to a surprising and novel result that the PLK inhibitor TPKI-30 targets also PYK2, currently an understudied kinase, and with a somewhat lesser potency also its paralog, FAK (PTK2, Fig. 6b).

Although previous work has demonstrated the potential of ML algorithms to help fill in the gaps in compound-target interaction maps,^16^ and to accelerate several phases of drug discovery,^21^ to date there has been no systematic and unbiased evaluations applied to comprehensive datasets. Participants of the Challenge made use of various ML modelling approaches, and rather surprisingly, no particular method class, training data source or bioactivity type stood out. Rather, the top-performing teams used relatively different approaches (Table 1). Some of the top-performing models used protein sequence as target feature, but no structural information. Furthermore, none of the top-performing models required 3D or other detailed chemical ginformation, making the ML models rather straightforward to apply for various compound and target classes. Recently, many advanced deep learning (DL) algorithms have been proposed for compound-target interaction prediction,^22–24^ but our results did not find DL outperforming other learning approaches. Interestingly, the Spearman correlation sub-challenge top-performer (Q.E.D) used the same modelling approach as the baseline model,^16^ yet showed markedly better performance (Fig. 3f), indicating that careful feature selection, method implementation, or other domain knowledge could result in marked performance improvement.

To get a more global picture, at the end of the Challenge we asked all the teams to fill in survey questionnaires to explore whether there would be any broad method classes or chemical or target features shared among the models. Among the 31 teams that answered the surveys, none of the method classes had a very strong contribution to the accuracy (Suppl. Fig. 11), similarly as has been seen also in other DREAM challenges.^25–27^ A rather surprising observation from the survey was that the K_d_ prediction accuracies could be improved by using other types of multi-dose bioactivity data (e.g. K_i,_ IC_50_, EC_50_), compared to using K_d_ data alone (Suppl. Fig. 11). This provides a further opportunity for ML models that often require relatively large training datasets, as these bioactivity types are among the most common in multi-dose target profiling, and more common than K_d_ in DTC database (Suppl. Fig. 11g). Another observation was that the teams that used DTC alone as training bioactivity data source had decreased predictive performance, perhaps due to the more heterogeneous bioactivity data stored in DTC, compared to BindingDB^11^ or ChEMBL.^10^ This suggests that further annotation and harmonization of the various types and sources of bioactivity data will be needed to make the most of these data for predictive modelling, ideally in the form of a crowdsourced community effort.

Many previous DREAM Challenges have demonstrated that ‘wisdom of the crowds’ may also improve the predictive power of the individual models through combining models as meta-predictors or ensemble models.^25–27^ The ensemble model constructed in this Challenge showed that the critical point came rather quickly after which adding more models led to rapid decrease in accuracy (Fig. 4d). The combination of the top-performing ML models improved both the sensitivity and specificity, compared to single-dose target activity assays, without requiring any additional experiments (Fig. 5). Notably, none of the top-performing models used single-dose inhibition data, and we showed how combining the inhibition measurements with ML models led to even higher prediction accuracy than using either one alone, while identifying an increased number of potent compound-kinase activities compared to when using the standard 80% inhibition cut-off (Fig. 6). Furthermore, the best-performing models were not dependent on the number or type of available bioactivity data, provided the training data had sufficient structural diversity for the kinase families being predicted. Subsequent experiments carried out based on the top-performing model predictions demonstrated that these models can facilitate experimental mapping efforts, both for well-studied and under-studied kinases (Fig. 6b,c).

To enable the community to apply the predictive models benchmarked in the Challenge to various drug development applications, we have made available the top-performing models as containerized source code. The Docker models enable continuous validation of the model predictions whenever new experimental kinase profiling data will become available, as well as make it possible to run the best performing models on private data that would otherwise remain closed and unavailable to the research community.^28^ This Challenge will, therefore, contribute to the further development and benchmarking of current and future target activity prediction models on a larger scale, possibly for other target classes. The systematically validated models can guide many precision medicine applications, such as prediction of selective inhibitors for new disease targets, or off-target potency predictions for investigational compounds. All the models, new bioactivity data, and benchmarking infrastructure are openly available on Synapse platform www.doi.org/10.7303/syn15667962) and DTC platform (https://drugtargetcommons.fimm.fi/). We envision that the IDG-DREAM Challenge will provide a continuously-updated resource for the chemical biology community to prioritize and experimentally test new target activities toward accelerating many drug discovery and repurposing applications.

## Online Methods

### Challenge infrastructure and timeline

The Challenge was organized and run on the collaborative science platform Synapse. All prediction files were submitted using the Challenge feature of this platform to track which teams and individuals submitted files, and to track the number of submissions per team. Challenge infrastructure scripts including code for calculating the scoring metrics are available at https://github.com/Sage-Bionetworks/IDG-DREAM-Drug-Kinase-Challenge. Teams were permitted to submit three predictions for Round 1, and two predictions for Round 2 (Suppl. Fig. 3). For Rounds 1 and 2, we used a common workflow language-based challenge infrastructure to perform the following tasks: (1) validate a prediction file to ensure that it conformed to the correct file structure and had numeric pK_d_ predictions and return an error email to participants if invalid, (2) run a python script to calculate the performance metrics for a submitted prediction, and (3) return the score to the Synapse platform. For Round 1b, in which we permitted 1 submission a day for 60 days, we implemented a modified Ladderboot^29^ protocol to prevent model overfitting. This was done by modifying step (2) above as follows: the scoring infrastructure receive a submitted prediction, check for a previous submission from the same team, and run an R script to bootstrap the current and previous submission 10,000 times, calculate a Bayes factor (K) between the two submissions; the scoring harness would then only return an updated score if it was substantially better (K > 3) than the previous submission.

### New bioactivity data for model testing

To generate unpublished test bioactivity data for scoring of predictions, we sent kinase inhibitors to DiscoverX (Eurofins Corporation) for the generation of new dose-response dissociation constant (K_d_) values, as a measure of a binding affinity. In order to give a better sense of the relative compound potencies, K_d_ is represented in the logarithmic scale, as pK_d_ = −log_10_(K_d_), where K_d_ is given in molars [M]. The higher the pK_d_ value, the higher the inhibitory ability of a compound against a protein kinase. The 95 inhibitors used in the Challenge (70 for Round 1 and 25 for Round were a part of the kinase inhibitor collection at the SGC-UNC for which we already had the single-dose inhibition screening done at DiscoverX across their large kinase panel. This scanMax^SM^ data (also called KINOMEScan) was collected at a screening concentration of 1 µM. A two-step screening approach was adopted, as in previous studies^4–6^, using the DiscoverX KINOMEscan standard protocol (https://www.discoverx.com/services/drug-discovery-development-services/kinase-profiling/kinomescan). The dose-response K_d_ values were generated for a range of compound-kinase pairs that had inhibition>80% in the single-dose assay. The compounds were supplied as 10 mM stocks in DMSO, and the top screening concentration was 10 mM.

A total of 25 of the axitinib-kinase pairs generated for Round 2 were already profiled in previous published studies,^6,15^ and were therefore excluded from the Round 2 test dataset. The Spearman correlation between these newly-measured pK_d_’s and those available from DTC was 0.701 (Suppl. Fig. 12a), providing the experimental consistency of the K_d_ measurements for axitinib. We note this 25 pK_d_’s is a rather limited set for such analysis of consistency, and therefore we extracted a larger set of 416 K_d_ measurements that overlapped with the Round 2 kinases from two comprehensive target profiling studies,^4,5^ including 104 pairs where pK_d_ = 5 in both of the studies. The Spearman correlation of these replicate pK_d_ measurements was 0.842 (Suppl. Fig. 12b), demonstrating a good reproducibility of the pK_d_ measurements. These replicate measurements were used when determining a practical upper limit for the predictive accuracy of the machine learning models in the scoring of their predictions (see below).

To subsequently test the top-performing model predictions in additional compound-kinase pairs that were not part of Round 1 or 2 datasets, we selected a set of 88 pairs that showed most potency based on the average predicted pK_d_ of the top-performing models (Q.E.D., DMIS-DK and AIWIC), regardless of their single-dose inhibition levels. These 88 pairs were actually scattered across the whole spectrum of single-dose inhibition levels, ranging from 0% to 78% (Supplementary Fig. 10; note: pairs with inhibition >80% were K_d_-profiled already in Round 2). One of the compounds (TPKI-35) was not available from IDG, so the predicted 7 kinase targets for that compound could not be tested experimentally, resulting in a dataset of total of 81 compound-kinase pairs that were shipped to DiscoverX for multi-dose K_d_ profiling. One of the compounds (GW819776) was shipped separately in a tube, whereas the other 14 compounds were supplied as 10 mM stocks in DMSO, and the K_d_ profiling done was done using the same KINOMEscan competitive binding assay protocol as for the Round 1 and Round 2 pairs.

### Scoring of the model predictions

We used the following six metrics to score the predictions from the participants:

- Root-mean-square error (RMSE): square root of the average squared difference between the predicted pK_d_ and measured pK_d_, to score continuous activity predictions.
- Pearson correlation: Pearson correlation coefficient between the predicted and measured pK_d_’s, which quantifies the linear relationship between the activity values.
- Spearman correlation: Spearman’s rank correlation coefficient between the predicted and measured pK_d_’s, which quantifies the ability to rank pairs in correct order.
- Concordance index (CI)^30^: probability that the predictions for two randomly drawn compound-kinase pairs with different pK_d_ values are in the correct order.
- F1 score: the harmonic mean of the precision and recall metrics. Interactions were binarized by their pK_d_ values into positive class (pK_d_ > 7) and negative class (pK_d_ ≤ 7).
- Average AUC: average area under ten receiver operating characteristic (ROC) curves generated using ten interaction threshold values from the pK_d_ interval [6, 8] to binarize pK_d_’s into true class labels.

The submissions in Round 1 were scored across the six metrics but the teams remained unranked. The Round 2 consisted of two sub-challenges, the top-performers of which were determined based on RMSE and Spearman correlation, respectively. Spearman correlation evaluated the predictions in terms of accuracy at ranking of the compound-kinase pairs according to the measured K_d_ values, whereas RMSE considers the absolute errors in the quantitative binding affinity predictions. The tie-breaking metric for both Rounds was averaged area under the curve (AUC) metric in the ROC analyses that evaluated the accuracy of the models to classify the pK_d_ values into active and inactive classes based on multiple K_d_ cutoffs.

### Statistical evaluation of the predictions

Determination of the top-performers was made by calculation of a Bayes factor relative to the top-ranked submission in each category. Briefly, we bootstrapped all submissions (10,000 iterations of sampling with replacement), and calculated RMSE and Spearman correlation to the test dataset to generate a distribution of scores for each submission. A Bayes factor was then calculated using the challengescoring R package (https://github.com/sage-bionetworks/challengescoring) for each submission relative to the top submission in each subchallenge. Submissions with a Bayes factor ≤ 3 relative to the top submission were considered to be tied as top-performers. Tie breaking for both subchallenges was performed by identifying submission with the highest absolute average AUC.

To create a distribution of random predictions, we randomly shuffled the 430/394 K_d_ values across the set of 430/394 compound-kinase pairs in the Round 1/Round 2 datasets, and repeated the permutation procedure 10,000 times. Then we compared the actual Round 1/Round 2 prediction scores to Spearman and RMSE calculated from the permuted K_d_ data. We defined a prediction as better than random if its score was higher than the maximum of the 10,000 random predictions (empirical *P* = 0.0, permutation test).

To determine the maximum possible performance practically achievable by any computational models, we utilized replicate K_d_ measurements from distinct studies that applied a similar biochemical assay protocol. We used the DrugTargetCommons to retrieve 863 and 835 replicated K_d_ values for kinases or compounds that overlapped with the Round 1 and 2 datasets, respectively. These data originated from two comprehensive screening studies^4,5^. To better represent the distribution of pK_d_ values in the test data, we subset the DTC data to contain 35% (Round 1) and 25% (Round 2) pK_d_=5 values, approximately matching the proportion of pK_d_ = 5 values in R1 and R2 test sets. For Round 1, we used 317 replicated K_d_s, including 111 randomly selected pairs where pK_d_ = 5. For Round 2, we used 416 replicated Kds, including 104 randomly selected pairs where pK_d_ = 5. We randomly sampled the replicate measurements of these compound-kinase pairs (with replacement), calculated the Spearman correlation and RMSE between the pKd’s of the two studies for each 430 and 394 sub-sampled sets for Round 1 and 2, respectively, and repeated this procedure for a total of 10,000 samplings.

### The baseline prediction model

We used a recently-published and experimentally-validated kernel regression framework as a baseline model for compound-kinase binding affinity prediction^16^. Our training dataset consisted of 44,186 pK_d_ values (between 1968 compounds and 423 human kinases) extracted from DTC. Median was taken if multiple pK_d_ measurements were available for the same compound-kinase pair. We constructed protein kinase kernel using normalized Smith-Waterman alignment scores between full amino acid sequences, and four Tanimoto compound kernels based on the following fingerprints implemented in rcdk R package^31^: (i) 881-bit fingerprint defined by PubChem (*pubchem*), (ii) path-based 1024-bit fingerprint (*standard*), (iii) 1024-bit fingerprint based on the shortest paths between atoms taking into account ring systems and charges (*shortestpath*), and (iv) extended connectivity 1024-bit fingerprint with a maximum diameter set to 6 (ECFP6; *circular*). We used CGKronRLS as a learning algorithm^32^ (implementation available at https://github.com/aatapa/RLScore). We conducted a nested cross-validation in order to evaluate the generalisation performance of CGKronRLS with each pair of kinase and compound kernels as well as to tune the regularisation hyperparameter of the model. In particular, since the majority of the compounds from the Challenge test datasets had no bioactivity data available in the public domain, we implemented a nested leave-compound-out cross-validation to resemble the setting of the Challenge as closely as possible. The model comprising of protein kernel coupled with compound kernel built upon path-based fingerprint (*standard*) achieved the highest predictive performance on the training dataset (as measured by RMSE), and therefore it was used as a baseline model for compound-kinase binding affinity prediction in both Challenge Rounds.

### Top-performing models

Supplementary write ups provide details of all qualified models submitted to the Challenge (http://www.doi.org/10.7303/syn21445941.1). The key components of the top-performing models are listed in Table 1 and summarized below.

#### Team Q.E.D model

To enable a fine-grained discrimination of binding affinities between similar targets (*e.g*., kinase family members), the team Q.E.D explicitly introduced similarity matrices of compounds and targets as input features into their regression model. The regression model was implemented as an ensemble version (uniformly averaged predictor) of 440 CGKronRLS regressors^32,33^, but with different choices of regularization strengths [0.1, 0.5, 1.0, 1.5, 2.0], training epochs [400, 410, …, 500], and similarity matrices: the protein similarity matrix was derived based on the normalized striped Smith-Waterman alignment scores^34^ between full protein sequences (https://github.com/mengyao/Complete-Striped-Smith-Waterman-Library). Eight different alternatives of compound similarity matrices were computed using both Tanimoto and Dice similarity metrics for different variants of 1024-bit Morgan fingerprints^35^(‘radius’ [2, 3] and ‘useChirality’ [True, False], implementation available at https://github.com/rdkit/rdkit). Unlike the baseline method, which used only the available pK_d_ values from DTC for training, the team Q.E.D model extracted 16945 pK_d_, 53894 pK_i_ and 3301 pEC_50_ values from DTC. After merging the same compound-kinase pairs from different studies by computing their medians, 60462 affinity values between 13608 compounds and 527 kinases were used as the training data.

#### Team DMIS_DK model

Team DMIS_DK built a multi-task Graph Convolutional Network (GCN) model based on 953521 bioactivity values between 474875 compounds and 1474 proteins extracted from DTC and BindingDB. Three types of bioactivities were considered, that is, pK_d_, pK_i_, and pIC_50_. Median was computed if multiple bioactivities were present for the same compound-protein pair. Multi-task GCN model was designed to take compound SMILES strings as an input, which were then converted to molecular graphs using RDKit python library (http://www.rdkit.org). Each node (i.e. atom) in a molecular graph was represented by a 78-dimensional feature vector, including the information of atom symbol, implicit valence, aromaticity, number of bonded neighbors in the graph, and hydrogen count. No protein descriptors were utilized. The final model was an ensemble of four multi-task GCN architectures described in the Supplementary writeups (http://www.doi.org/10.7303/syn21445941.1). For the Challenge submission, the binding affinity predictions from the last K epochs were averaged, and then the average was taken over the 12 multi-task GCN models (four different architectures with three different weight initializations). Hyper-parameters of the multi-task GCN models were selected based on the performance on a hold-out set extracted from the training data. The GCN models were implemented using PyTorch Geometric (PyG) library^36^.

#### Team AI Winter is Coming model

Team AI Winter is Coming built their prediction model using Gradient Boosted Decision Trees (GBDT) implemented in XGBoost algorithm^37^. Training dataset included 600000 pK_d_, pK_i_, pIC_50_, and pEC_50_ values extracted from DTC and ChEMBL (version 25), considering only compound-protein pairs with ChEMBL confidence score of 6 or greater for ‘binding’ or ‘functional’ human kinase protein assays. For a given protein target, replicate compounds with different bioactivities in a given assay (differences larger than one unit on a log scale) were excluded. For similar replicate measurements, a single representative assay value was selected for inclusion in the training dataset. Each compound was characterized by a 16000-dimensional feature vector being a concatenation of four ECFP fingerprints (as implemented in RDKit) with a length set to of 5, 7, 9, and 11. No protein descriptors were used in the XGBoost algorithm^37^. A separate model for each protein target was trained using nested cross-validation (CV), where inner loops were used to perform hyperparameter optimisation and recursive feature elimination. The final binding affinity prediction was calculated as an average of the predictions from the cross-validated models based on five outer CV loops.

### Mean ensemble model construction

Ensemble models were generated by combining the best-scoring Round 2 predictions from each team. We iteratively combined models starting from the highest scoring Round 2 prediction (e.g. ensemble #1 - highest scoring prediction, ensemble #2 - 2 highest scoring, ensemble #3 - 3 highest scoring, and so on) for all 54 Round 2 submissions. Three types of ensembles were created using arithmetic mean, median, and rank-weighted summarization approaches. The rank-weighted ensemble was calculated by multiplying each set of predictions by the total number of submissions plus 1 minus the rank of the prediction file, summing these weighted predictions, and then dividing by the sum of the multiplication factors. The 54 ensemble predictions for each of the 3 summary metrics were bootstrapped and Bayes factors were calculated as previously described to determine which models were substantially different than the top ranked submission.

### Estimating the expected inhibition levels

The KINOMEscan assay protocol utilized for both the single-dose and dose-response assays is based on competitive binding assays, where the maximum compound concentration tested was 1 µM in both of the assays. For a given compound-kinase pair, the K_d_ values calculated from the dose-response assay were then used to estimate the expected single-dose %inhibition level (at 1 µM of compound) using the conventional ligand occupancy formula:

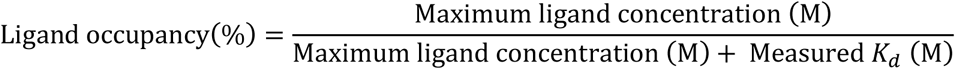

Here, the maximum ligand concentration is 10^−6^ M in the kinase assay. Therefore, a measured pK_d_ = 3 (i.e. K_d_ =10^−3^ M) results in the expected inhibition of 0%, pK_d_ = 4 and 5 in 1% and 10% expected inhibitions, respectively, and pK_d_ = 9 (i.e. K_d_ =10^−9^ M) results in expected inhibition of 100%.

#### Activity classification analyses

The standard confusion matrix was constructed using the measured pK_d_ values to define the true positive and true negative classes for the 394 pairs in Round2, using either pK_d_ > 6 and pK_d_ > 7 for indicating true positive activity. The predicted positive and negative classes for the pairs were defined based on either the single-dose activity measurement, using inhibition cut-off of 80%,^6,15,17^ or the model-predicted pK_d_ values, using the same activity thresholds as with the measured pK_d_ values (i.e., either pK_d_ = 6 or pK_d_ = 7). Positive predictive value (PPV) and false discovery rate (FDR) were calculated as the classification performance scores. The lower threshold of measured pK_d_ = 6 was used in the classification evaluations to have more balanced true positive and negative classes. To carry out a more systematic analysis of the model prediction accuracies, the 394 pairs in Round 2 were ranked both using the model-predicted pK_d_ values and the measured single-dose %inhibition values, and then these rankings were compared against the ground-truth activity classification based on the dose-response measurements (using again either pK_d_ > 6 and pK_d_ > 7 for indicating the true positive activity). The results were visualized using both receiver operating characteristic (ROC) and precision-recall (PR) curves, implemented in the pROC and pRROC R-packages, respectively^38,39^. The area under the ROC and PR curves was calculated as summary classification performance.

## Data and code availability

The Challenge test data will be made available in DTC (https://drugtargetcommons.fimm.fi/). The Docker containers of the best-performing teams are available on Synapse project (www.doi.org/10.7303/syn15667962). The codes for reproducing the results and figures are available at GitHub (https://github.com/Sage-Bionetworks/IDG-DREAM-Challenge-Analysis/). Key R-packages used beyond those mentioned elsewhere in Methods include tidyverse^40^ and the Synapse Python Client (https://github.com/Sage-Bionetworks/synapsePythonClient); all the packages used in the work and their versions can be found in the renv lockfile in the above GitHub repository.

### Acknowledgements

The authors thank the IDG Kinase Data and Resource Generation Center for generating new sets of target activity data for the Challenge Rounds 1 and 2, Olle Hansson (FIMM) for technical assistance with DrugTargetCommons platform, Tianduanyi Wang (FIMM) for his help with the baseline submissions, Anna Goldenberg (University of Toronto, Canada) and Chloe-Agathe Azencott (Institut Curie, France) for organizing the DREAM Idea Challenge, and Barbara Rieck and Ladan Naghavian for the bioactivity profiling at DiscoverX (Eurofins Corporation).

## IDG-DREAM Drug Kinase Binding Prediction Challenge Consortium members

**User oselot:** Mehmet Tan; **Team N121:** Chih-Han Huang, Edward S. C. Shih, Tsai-Min Chen, Chih-Hsun Wu, Wei-Quan Fang, Jhih-Yu Chen, and Ming-Jing Hwang; **Team Let_Data_Talk:** Xiaokang Wang, Marouen Ben Guebila, Behrouz Shamsaei, Sourav Singh; **User thinng:** Thin Nguyen; **Team KKT:** Mostafa Karimi, Di Wu, Zhangyang Wang, Yang Shen, **Team Boun:** Hakime Öztürk, Elif Ozkirimli, and Arzucan Özgür; **Team Aydin:** Zafer Aydin, Halil Ibrahim Bilgin; **Team KinaseHunter:** Hansaim Lim, Lei Xie; **Team AmsterdamUMC-KU-team:** Georgi K. Kanev, Albert J. Kooistra, Bart A. Westerman; **Team DruginaseLearning:** Panagiotis Terzopoulos, Konstantinos Ntagiantas, Christos Fotis, Leonidas Alexopoulos; **Team KERMIT-LAB - Ghent University:** Dimitri Boeckaerts, Michiel Stock, Bernard De Baets, Yves Briers; **Team QED:** Fangping Wan, Shuya Li, Yunan Luo, Hailin Hu, Jian Peng, Jianyang Zeng; **Team METU_EMBLEBI_CROssBAR:** Tunca Dogan, Ahmet S. Rifaioglu, Heval Atas, Rengul Cetin Atalay, Volkan Atalay, Maria J. Martin; **Team DMIS_DK:** Sungjoon Park, Minji Jeon, Sunkyu Kim, Junhyun Lee, Seongjun Yun, Bumsoo Kim, Buru Chang, Jaewoo Kang; **Team AI Winter is Coming:** Mariya Popova, Stephen Capuzzi, Olexandr Isayev; **Team hulab:** Gábor Turu, Ádám Misák, Bence Szalai, László Hunyady; **Team ML-Med:** Matthias Lienhard, Paul Prasse, Ivo Bachmann, Julia Ganzlin, Gal Barel, and Ralf Herwig, **Team Prospectors:** Davor Oršolic, Bono Lucic, Višnja Stepanic, Tomislav Šmuc; **Challenge organizers:** Anna Cichonska, Balaguru Ravikumar, Robert J Allaway, Michael Mason, Andrew Lamb, Zia-ur-Rehman Tanoli, Kristen Dang, Tudor I. Oprea, Avner Schlessinger, David H. Drewry, Gustavo Stolovitzky, Krister Wennerberg, Justin Guinney, Tero Aittokallio

