## Supplemental Figures for "Crowdsourced mapping extends the target space of kinase inhibitors"

Supplementary Figures 1-12

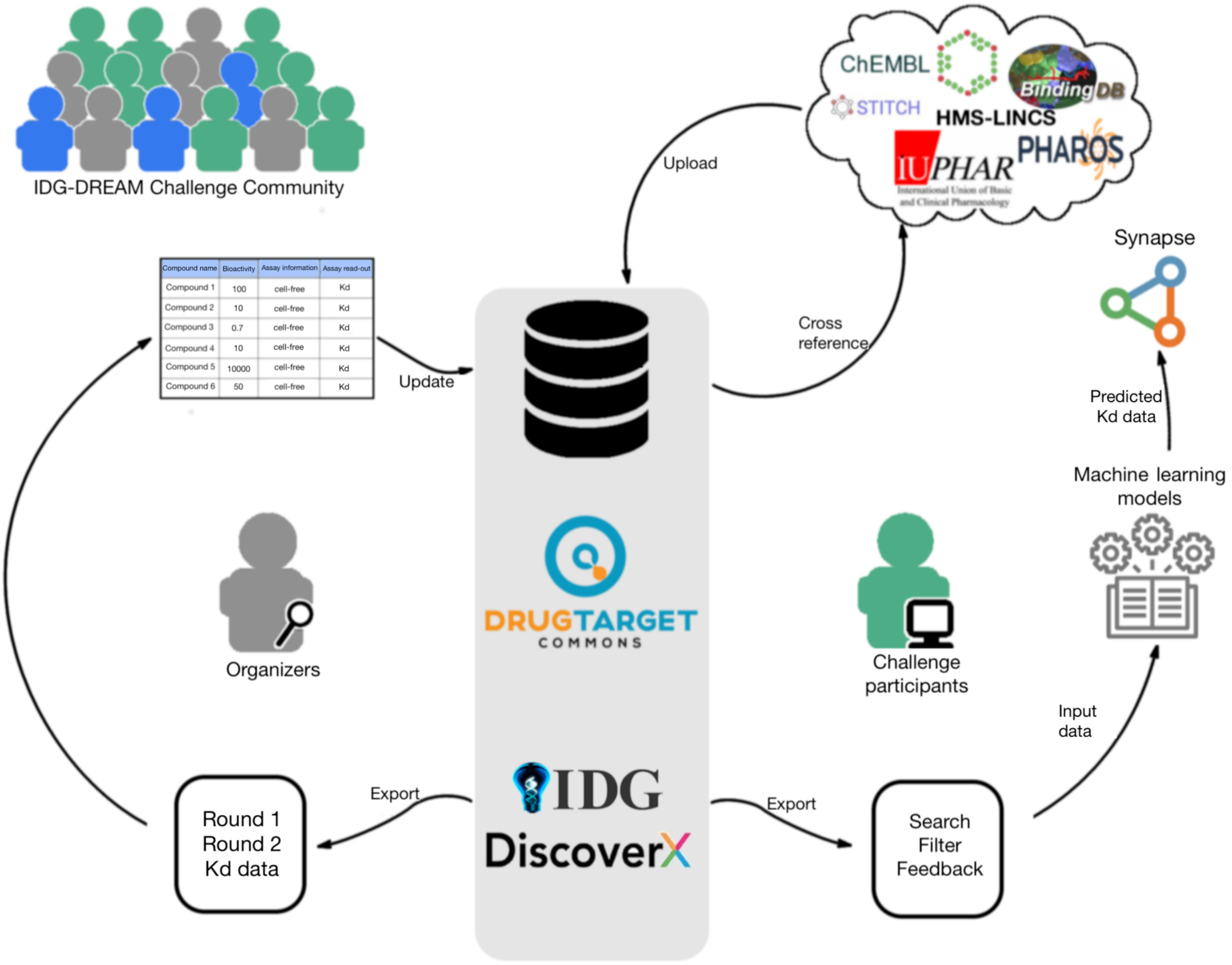


**Supplementary Figure 1**. The use of DrugTargetCommons (DTC) open-data platform in the Challenge. The test bioactivity data were provided by the Illuminating the Druggable Genome (IDG) program, and the multi-dose dissociation constant ($K_d$) data for Rounds 1 and 2 were generated by DiscoverX (Eurofins Corporation).

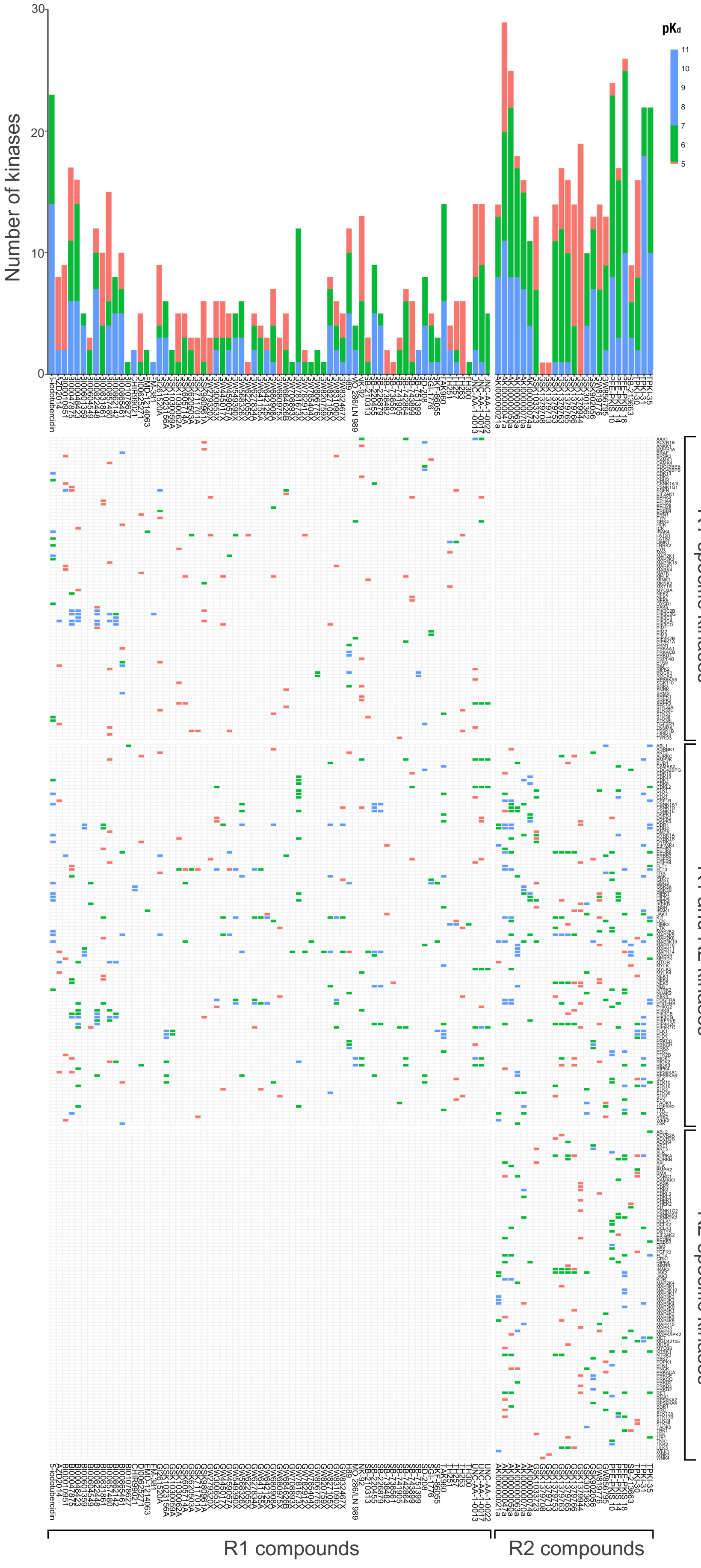


**Supplementary Figure 2.** Multi-dose dissociation constant ($K_d$) heatmap and distributions of the compound-kinase activities used as the test data in the Challenge Rounds 1 and 2.

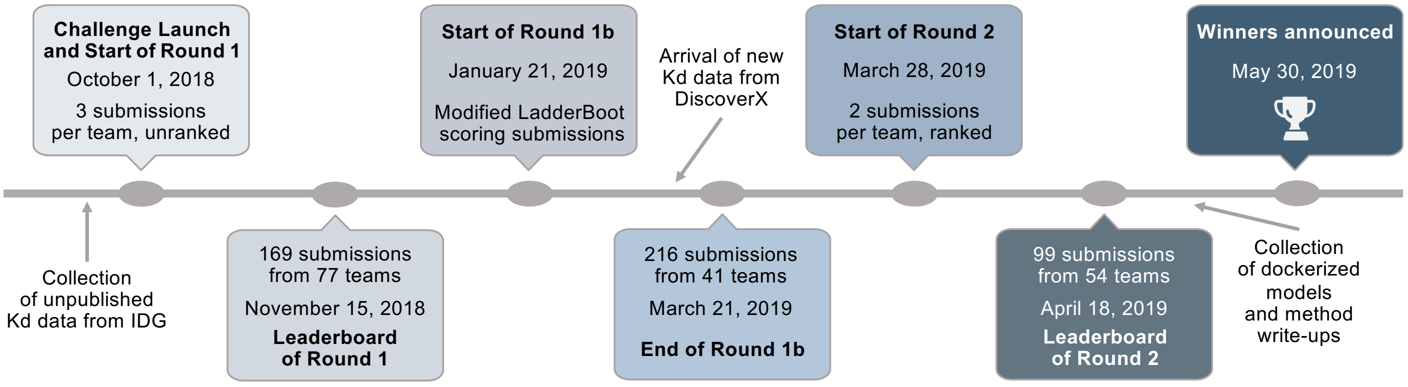


**Supplementary Figure 3.** Timeline of the IDG-DREAM Drug-Kinase Binding prediction Challenge. The ad-hoc leaderboard Round 1b was implemented while waiting for the new dissociation constant ($K_d$) bioactivity data from DiscoverX.

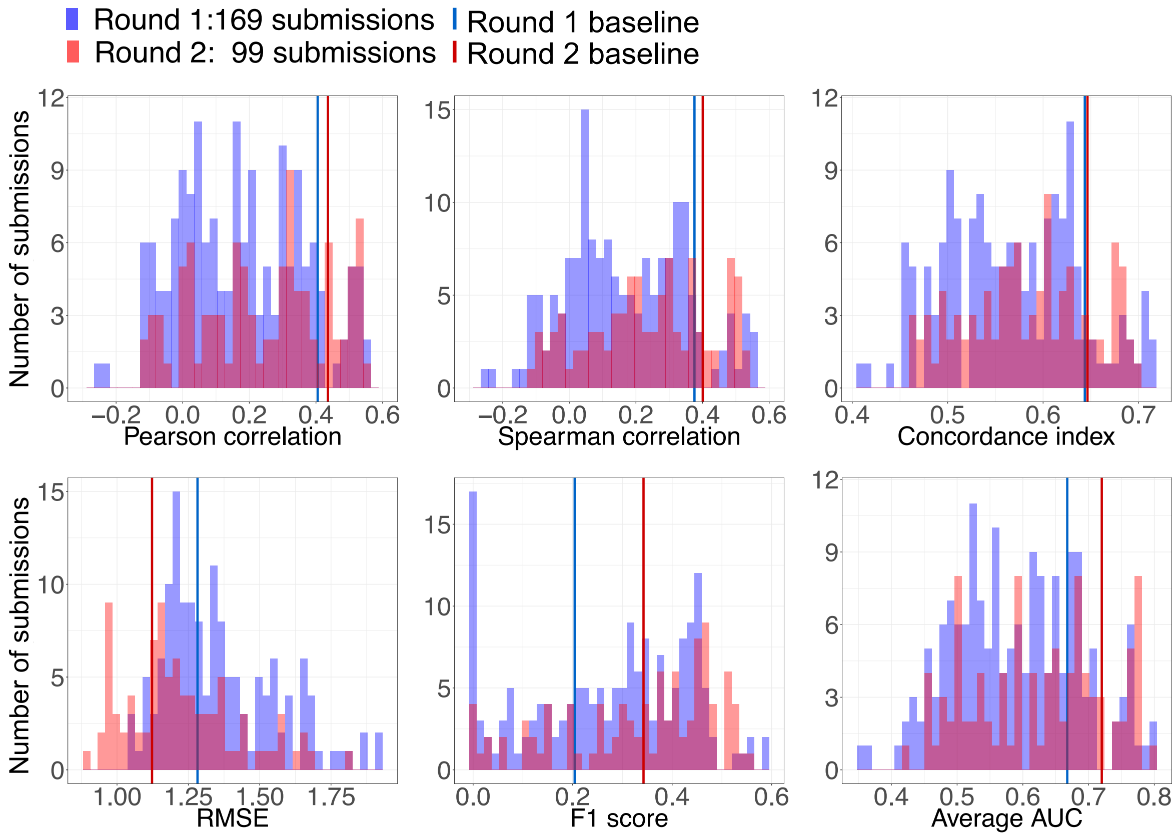


**Supplementary Figure 4.** Distributions of the Round 1 and Round 2 predictions, as evaluated with six scoring metrics, and compared to the baseline model (horizontal lines).

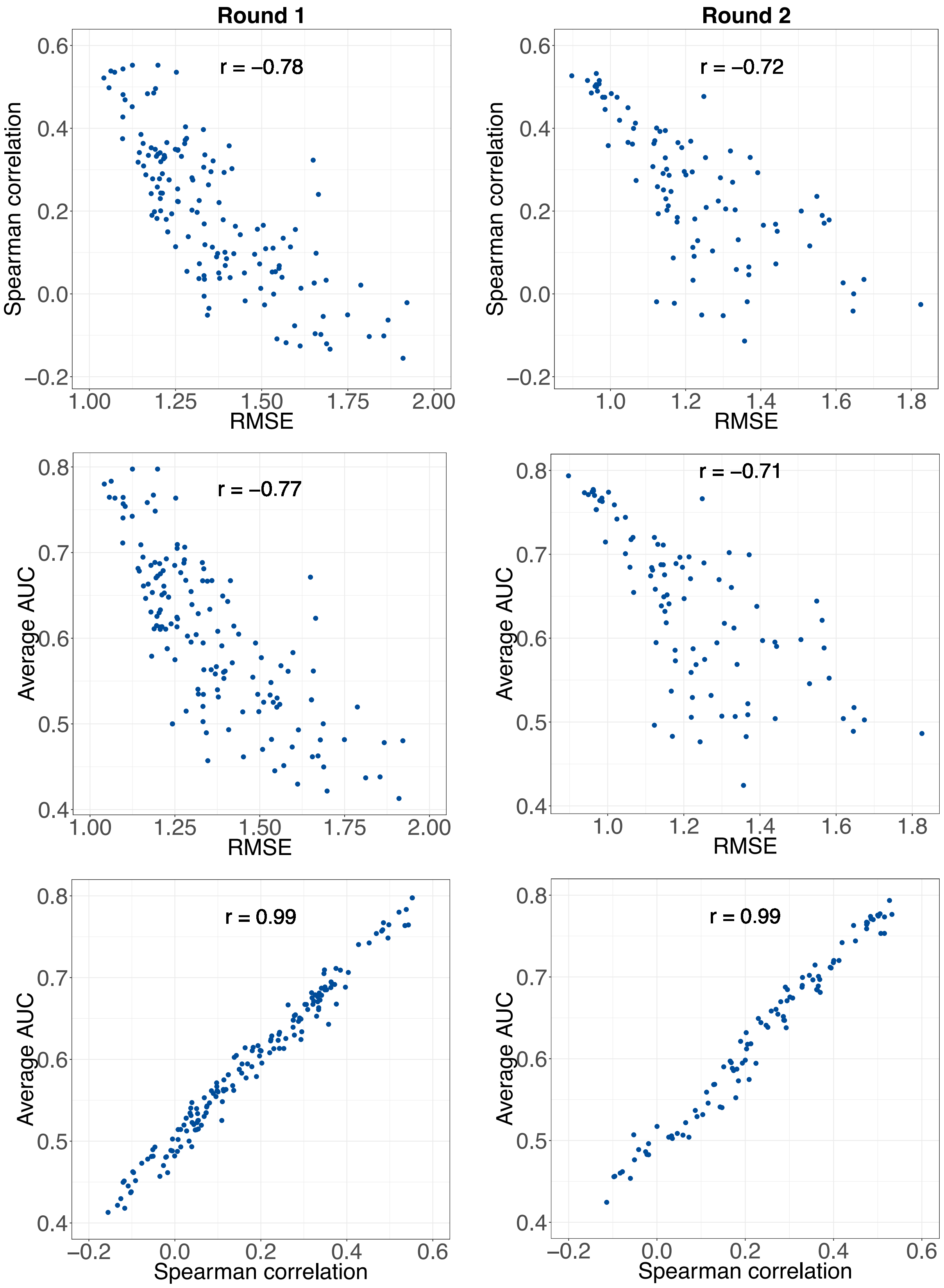


**Supplementary Figure 5.** Relationship between the two winning metrics (Spearman correlation and RMSE), and the tie-breaking metric (average AUC). Each point corresponds to one of the 169 submissions in Round 1 (left panel), or 99 submissions in Round 2 (right panel).

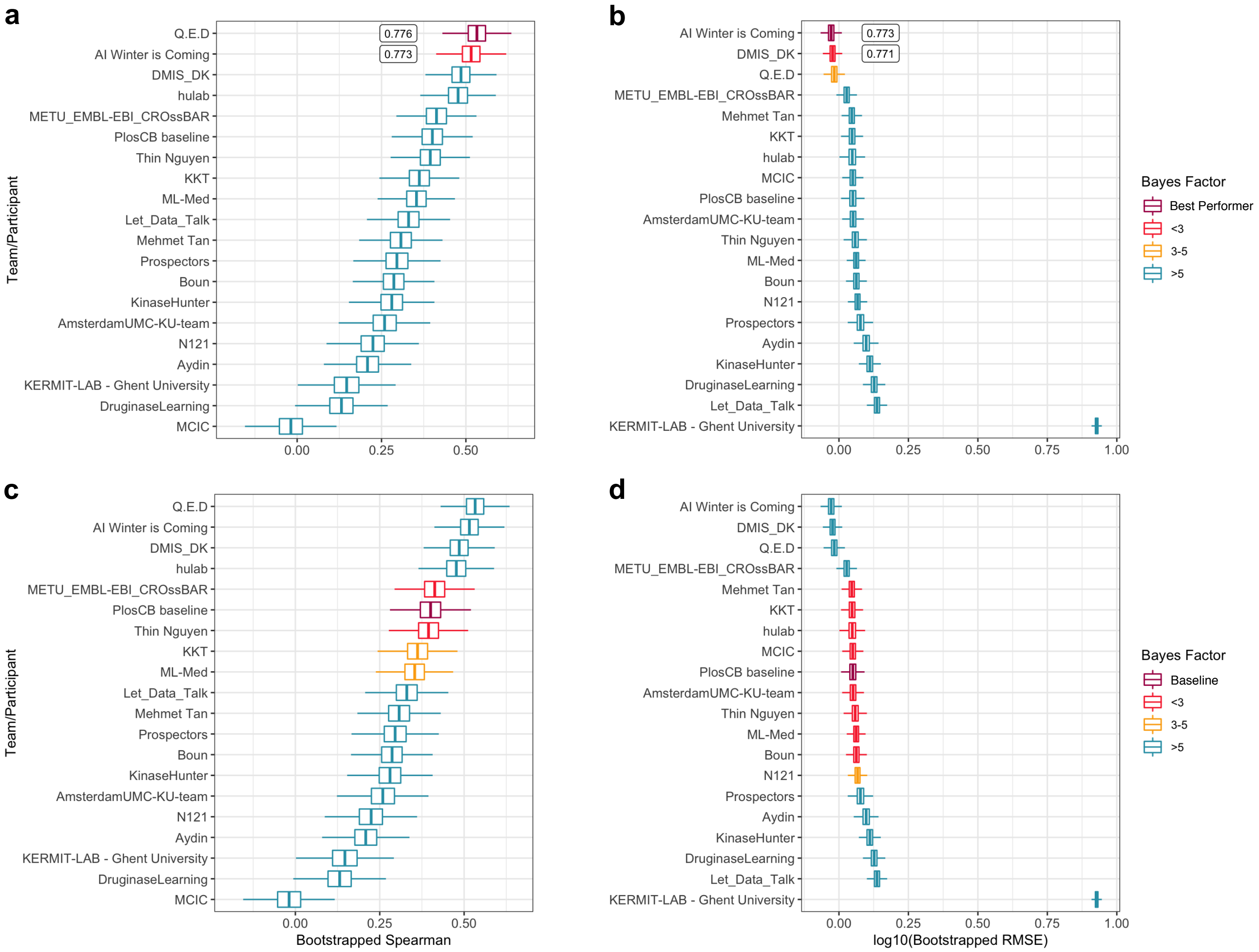


**Supplementary Figure 6**. Bayes factor analysis of the qualified participants/teams in Round 2. Upper panel: comparison against the top-performing model using bootstrapped Spearman correlation (a) and bootstrapped RMSE (b). Lower panel: comparison against the baseline model using bootstrapped Spearman correlation (c) and bootstrapped RMSE (d).

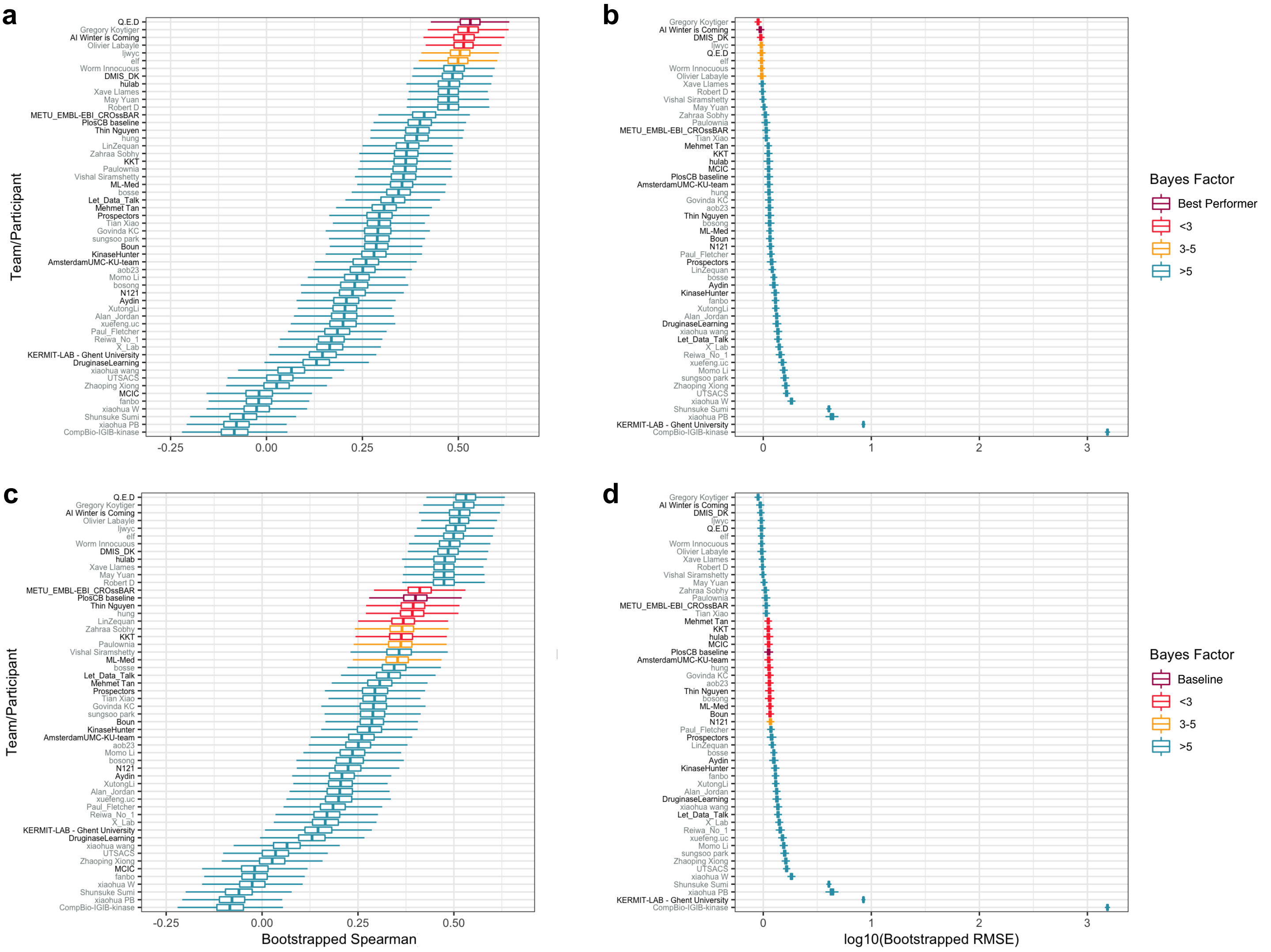


**Supplementary Figure 7**. Bayes factor analysis of all the participants/teams in Round 2 (qualified participant/team names written in black font). Upper panel: comparison against the top-performing model using bootstrapped Spearman correlation (a) and bootstrapped RMSE (b). Lower panel: comparison against the baseline model using bootstrapped Spearman correlation (c) and bootstrapped RMSE (d).

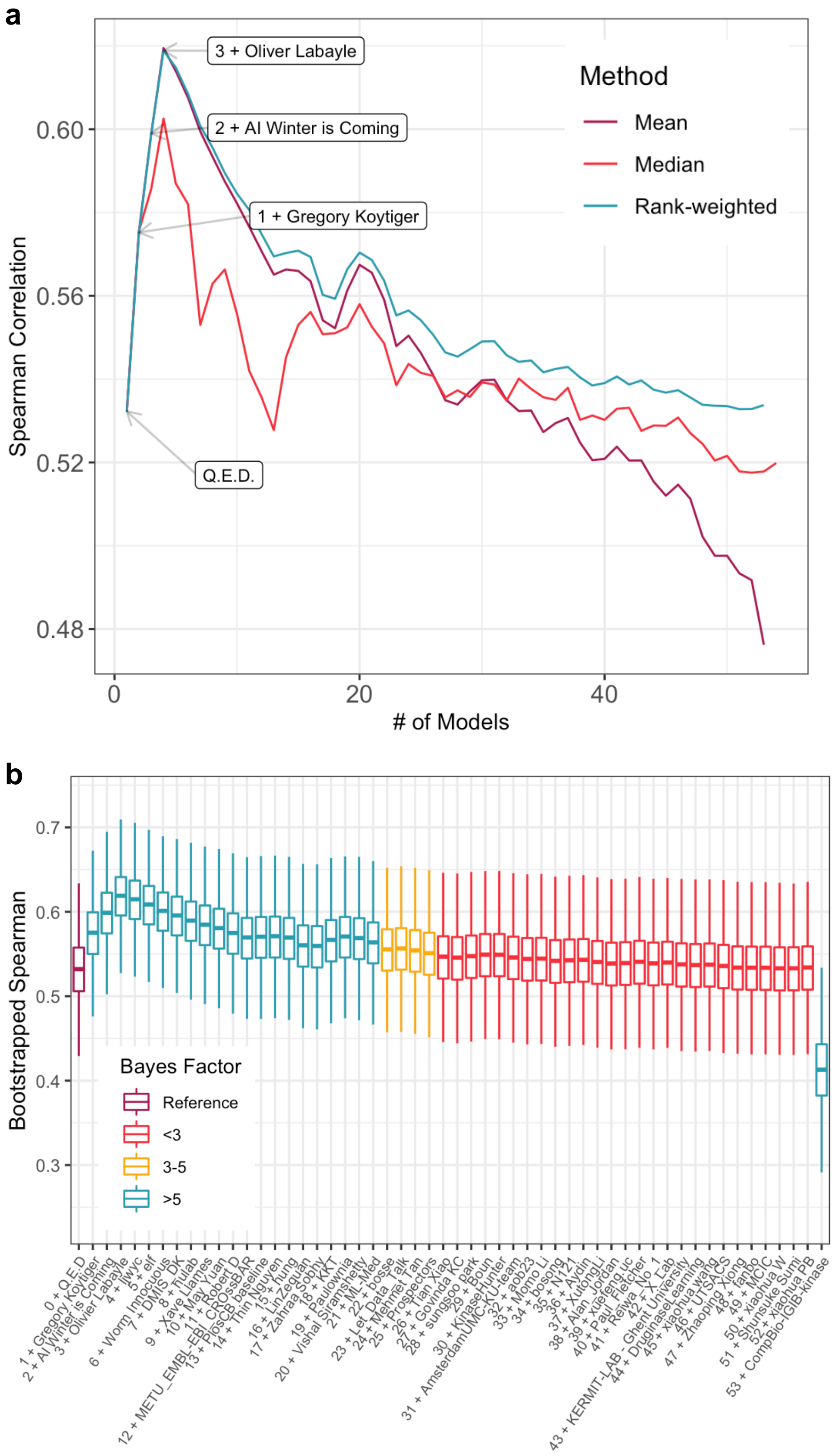


**Supplementary Figure 8.** Ensemble model construction. (a) Performance of the ensemble models when adding an increasing number of participants based on their Spearman correlation. Three different model aggregation methods were tested (the color trace legend). (b) Bayes factor analysis of the mean aggregation ensemble model compared to the top-performing reference (Q.E.D), based on bootstrapped Spearman correlation.

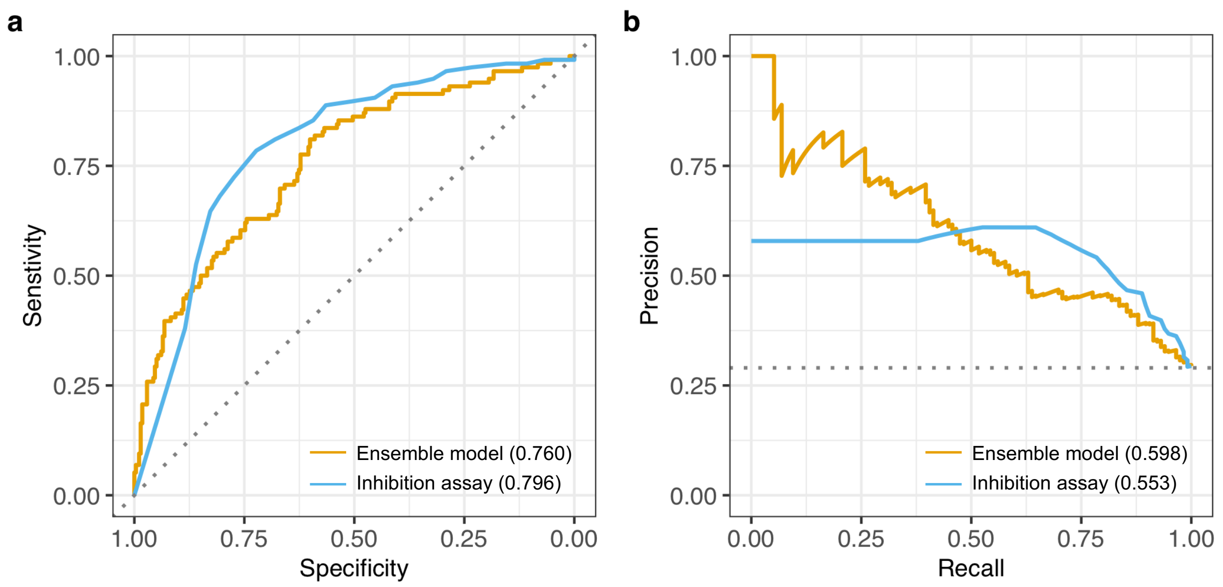


**Supplementary Figure 9.** (a) Receiver operating characteristic (ROC) curves when ranking the 394 pairs from Round 2 using the top-performing models' ensemble (average predicted $pK_d$) and the single-dose inhibition assays (the true positive activity class includes pairs with measured $pK_d>7$). The area under ROC curve values are shown in the parentheses and the diagonal dotted line shows the random prediction accuracy of AU-ROC=0.50. (b) Precision-recall (PR) curves for the same classification analysis than shown in panel a. The area under the PR curve values are shown in parentheses and the horizontal dotted line indicates the random classifier precision of 0.29.

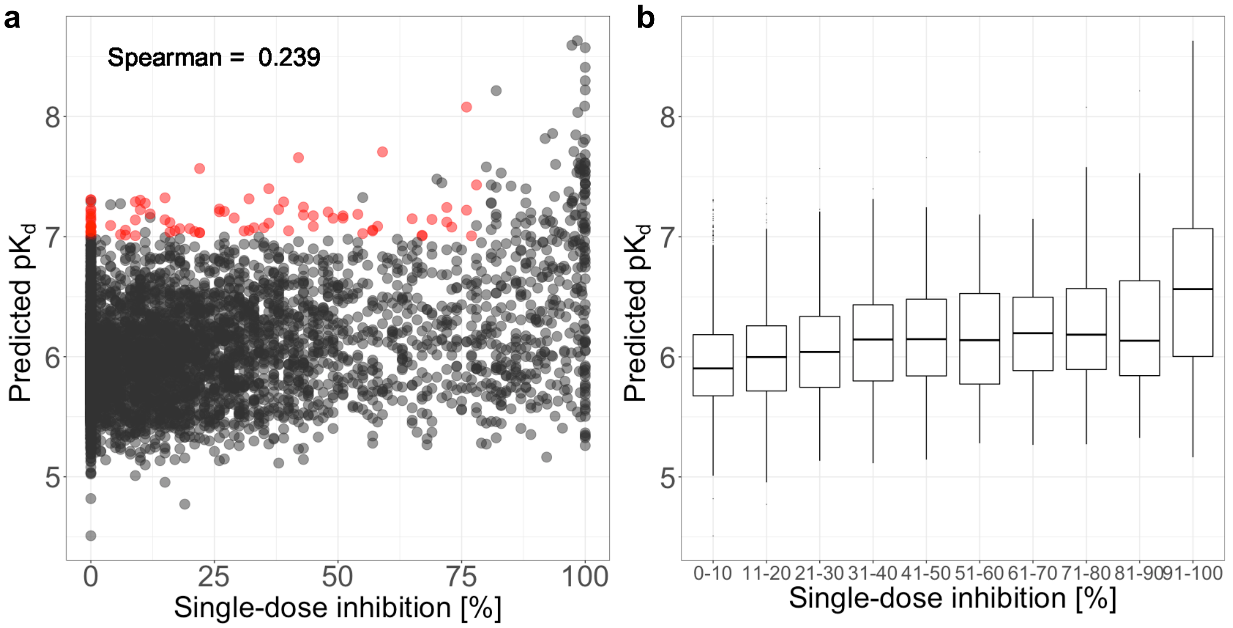


**Supplementary Figure 10.** (a) Average $pK_d$ prediction of the top-performing models' ensemble against single-dose inhibition levels in the full Round 2 data matrix, consisting of 5100 pairs between 25 inhibitors and 204 kinases that had %inhibition measurements available. Interestingly, the predictions models did not use any single-dose activity data in their training, and therefore showed only a marginal correlation with the measured %inhibition levels. (b) The same data analyzed using barplots to better show the relationships at the lowest (0%) and highest (100%) single-dose inhibition levels. The red points indicate potential false negatives based on the single-dose inhibition assay that have relatively high predicted $pK_d>7$ but were not $K_d$ profiled for the Round 2 dataset due to relatively low single-dose inhibition assay activity.

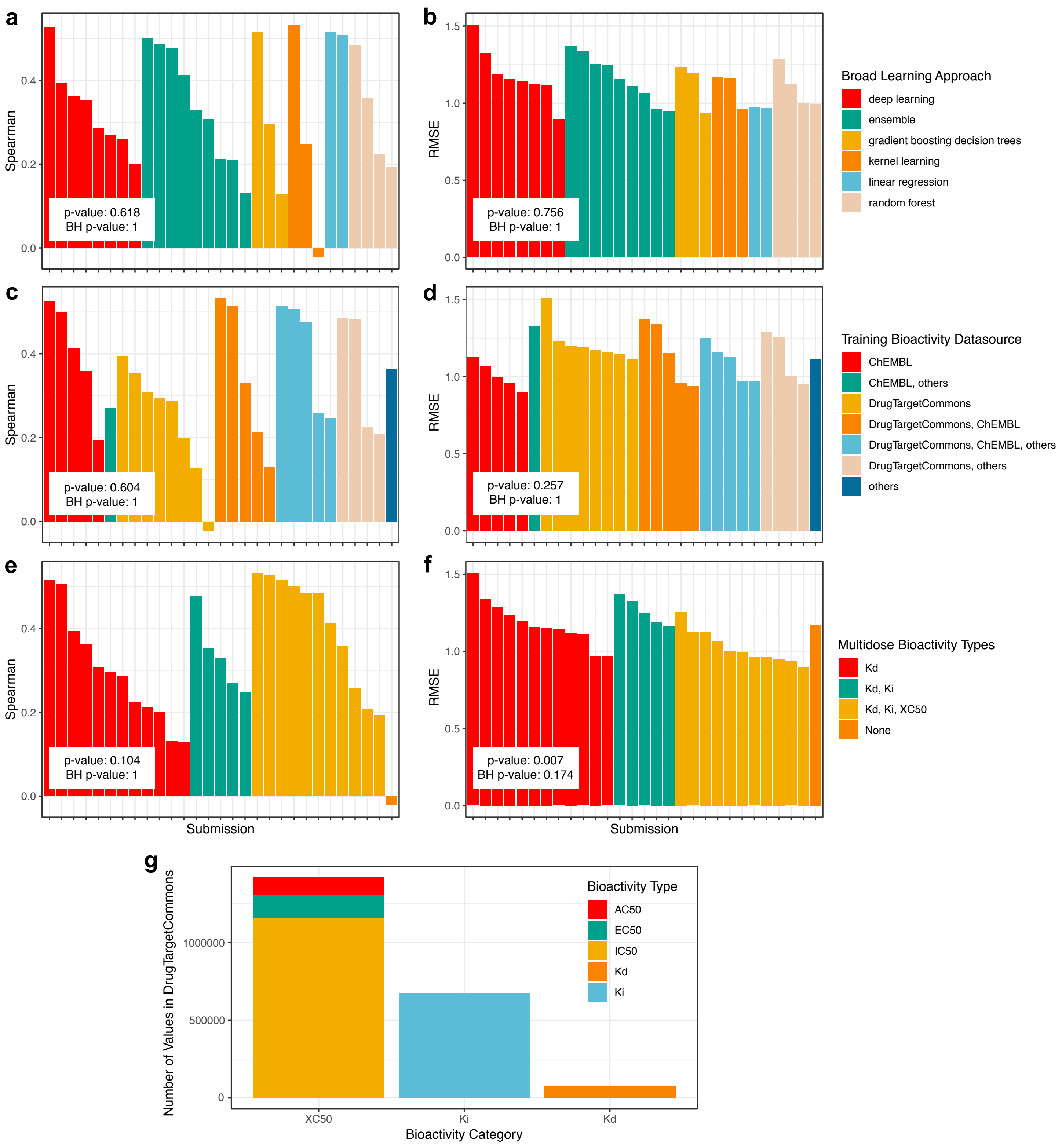


**Supplementary Figure 11.** The survey questionnaire results concerning the broad learning approaches, bioactivity data resources and multi-dose bioactivity datatypes used in model training. Statistical significance was assessed using the Kruskal-Wallis test (unadjusted p-values), and adjusted with Benjamini-Hochberg control of false discovery rate (FDR).

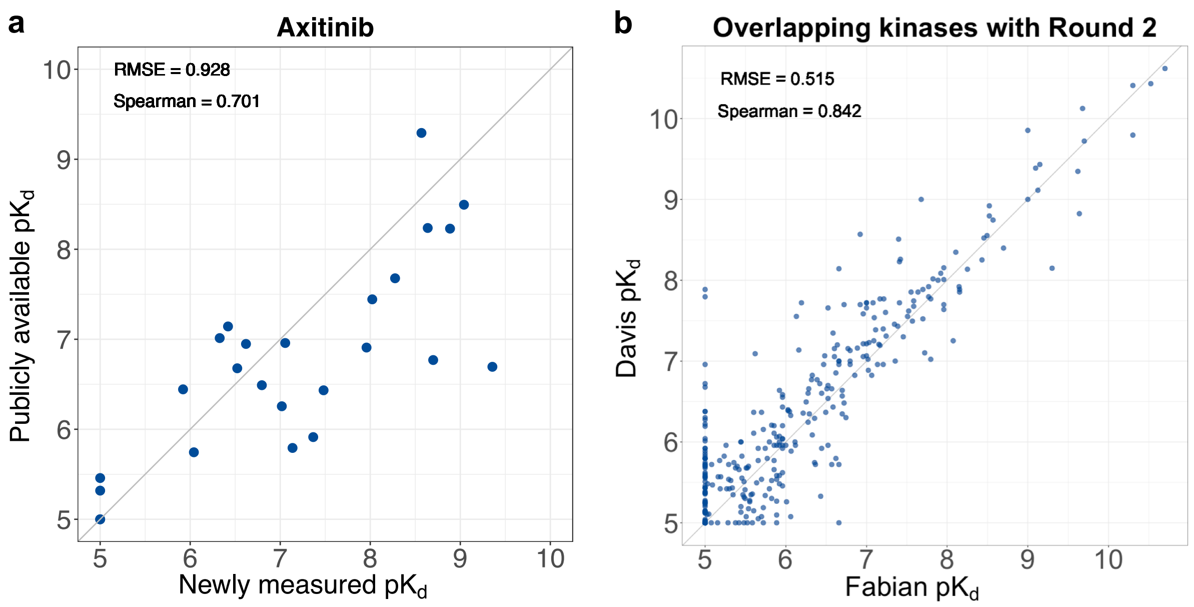


**Supplementary Figure 12.** (a) $pK_d$ values of the 25 axitinib-kinase pairs generated for Round 2 compared to those available in DTC. (b) $pK_d$ values of 412 compound-kinase pairs for kinases that overlapped with the Round 2 kinases from two comprehensive target profiling studies carried out by Davis et al. (PMID: 22037378) and Fabian et al. (PMID: 15711537). Out of the 412 pairs, 103 pairs have $pK_d$=5 in both of the studies, leading to relatively low RMSE.
